# Drug screening identifies the Src/Abl inhibitor Dasatinib as suppressor of IL-23 signalling in skin inflammation

**DOI:** 10.64898/2025.12.04.692333

**Authors:** MJ Valle-Pastor, I Cayuela, JS Yebra, L Senach-Rasilla, G Pastor-Fernández, A Oliva, J Traba, MN Navarro

## Abstract

IL-23-driven IL-17 production by γδ17 and Th17 cells is central to the pathogenesis of psoriasis and other autoimmune disorders. The intracellular mechanisms linking IL-23 signalling to effector cytokine production remain incompletely defined and thus, therapeutic targets downstream of IL-23 are limited. Here, we developed an *in vitro* model based on a γδ17 T cell line to study IL-23 responses and identify pharmacological inhibitors of type 3 immunity. We conducted a drug-repurposing screening of FDA-approved compounds and identified Src/Abl kinase inhibitors Dasatinib and Bosutinib as potent suppressors of IL-23-induced IL-17A production. In vivo, intraperitoneal and topical Dasatinib administration reduced epidermal thickening, immune cell infiltration, and the accumulation of IL-17-producing T cells in the Imiquimod model of skin inflammation. Mechanistically, Dasatinib and Bosutinib blocked IL-23-dependent mTORC1 and mTORC2 activation. Finally, loss-of-function experiments identified the Src kinase Blk as a critical mediator of IL-23-induced mTORC1 activation and IL-17A production in γδ17 T cells. These findings uncover a novel IL-23-Src-mTOR axis in type 3 immunity, opening the door for therapeutic targeting of Src signalling networks in IL-23/IL-17-driven inflammation.

## Introduction

Type 3 immunity is a specialized arm of the immune system primarily driven by interleukin 17 (IL-17) producing cells such as Th17 CD4 T cells, γδ17 T cells and innate lymphoid cells type 3 (ILC3s). Type 3 immune response plays a pivotal role in mucosal defence, targeting extracellular microbial threats and providing protective immunity against extracellular bacteria and fungi at mucosal barriers. While essential for host protection, dysregulated type 3 immunity is closely linked to the development of autoimmune and inflammatory diseases, including psoriasis, psoriatic arthritis, multiple sclerosis, ankylosing spondylitis or inflammatory bowel diseases (Annunziato, Romagnani and Romagnani, 2015; Mills, 2023).

Type 3 immune cells are characterised by the expression of the master transcription factor RORγt (encoded by *Rorc*), which is essential for IL-17 production and also promotes the expression of the interleukin 23 receptor (IL-23R) (Ivanov *et al*., 2006; Mezghiche *et al*., 2024). IL-23 is a key cytokine that drives the expansion, maintenance and activation of IL-17-producing T cells such as Th17 and γδ17 subsets, making the IL-23/IL-17 axis a validated therapeutic target in psoriasis and other disorders. In fact, neutralising antibodies targeting IL-23 and IL-17 have been approved for the treatment of severe plaque psoriasis. Despite their efficacy, antibody therapy prescription is mostly limited to severe symptoms. In contrast, small-molecule drugs offer greater flexibility: they can be administered orally, topically, and are capable of targeting intracellular proteins such as kinases and transcription factors. A deeper understanding of the signalling pathways that control IL-17 production downstream of IL-23 stimulation could enable the development of more precise therapeutic strategies to restore immune homeostasis in autoimmune and inflammatory conditions.

IL-23-driven cytokine production, primarily IL-17A but also IL-22, GM-CSF, and TNFα, plays a central pathogenic role in mouse models of psoriasis (Chan *et al*., 2006; Zheng *et al*., 2007; van der Fits *et al*., 2009; Hartwig *et al*., 2018). In this context, while Th17 cells contribute to disease pathology (van der Fits *et al*., 2009), γδ17 T cells play a dominant pathogenic role and they are major producers of IL-17A in the skin and other barrier tissues in mouse models (Cai *et al*., 2011; Pantelyushin *et al*., 2012; Yoshiki *et al*., 2014). γδ17 T cells have been also detected in skin lesions of psoriatic patients (Mills, 2023). γδ17 T cells are a subset of γδ (gamma-delta) T cells characterized by their ability to produce large amounts of IL-17A in response to immune challenges. γδ17 T cells commit to IL-17 production during thymic development (Fiala, Gomes and Silva-Santos, 2020; Inácio *et al*., 2025). Additionally, extrathymic signals such as IL-23 and IL-1β induce the differentiation of uncommitted γδ T cells into γδ17 effector T cells upon immune challenge (Muschaweckh, Petermann and Korn, 2017; Papotto *et al*., 2017).

The study of signalling pathways that regulate the pathogenic potential of γδ17 T cell and other IL-23 responders has been limited, largely due to the scarcity of these cell types. In this work, we have established an *in vitro* model based on a γδ17 T cell line to study IL-23 responses, and developed a drug-repurposing screening using 88 FDA-approved compounds to identify pharmacological inhibitors of type 3 immunity. We identified the dual Src/Abl inhibitors Dasatinib and Bosutinib as potent suppressors of IL-23-mediated IL-17 production. In vivo, both intraperitoneal and topical Dasatinib administration ameliorated different disease parameters in Imiquimod-induced model of skin inflammation. We found that Dasatinib and Bosutinib blocked the activation of mTORC1 and mTORC2 in response to IL-23 stimulation, signalling nodes previously shown to be required for IL-17 production (Cai *et al*., 2019; Cibrian *et al*., 2020). Furthermore, silencing the Src-famly kinase Blk recapitulated the lack of mTORC1 activation and IL-17 production in response to IL-23 stimulation. Together, these findings identify a previously unrecognized IL-23-Src-mTOR signalling axis in type 3 immunity, opening the door for therapeutic targeting of Src signalling networks in IL-23/IL-17-driven inflammation.

## Materials and methods

### Mice

IL-23R reporter mice with GFP knocked into the *Il23r* locus (*Il23r*-gfp) were kindly provided by Dr. F. Powrie (University of Oxford, UK), with the permission of Dr. M. Oukka (University of Washington, Seattle, USA) who originally generated this mouse strain (Awasthi *et al*., 2009). Heterozygous mice were used for this study (*Il23r*^wt/gfp^). C57BL/6J mice were obtained from Charles Rivers. Mice were maintained under specific pathogen–free (SPF) conditions at the animal facility of the Centro de Biología Molecular Severo Ochoa. Mice breeding and procedures were performed in accordance with national and institutional guidelines for animal care (EU Directive 2010/63/EU for the protection of animals used for scientific purposes). The experimental procedures were approved by the Director General de Medio Ambiente de la Comunidad de Madrid (Approval reference: PROEX 235/17 and PROEX010.0/23).

### Primary and Long Term IL-7 Cultures of γδ17 T cells (γδ17 T-LTICs)

Pooled LN from 5-10 mice (12-24 weeks) were mechanically disaggregated and γδT cells were isolated by positive magnetic sorting. Total LN cells were incubated with a biotin-coupled anti-TCRγδ antibody (GL3; BD Pharmingen), followed by incubation with Streptavidin Microbeads, and applied to LS magnetic columns (Miltenyi Bitotec). Isolated TCRγδ cells were cultured in home-made IMDM supplemented with 200mM L-glutamine, 10% (v/v) heat-inactivated fetal bovine serum (FBS; Biowest), penicillin (50 U/ml; Invitrogen), streptomycin (50 mg/ml; Invitrogen), and 2-mercaptoethanol (50 µM; Sigma) (IMDM-10%-2ME), supplemented with IL-7 (5 ng/ml; Miltenyi). For primary γδ17 T cell cultures, isolated TCRγδ cells were cultured in IL-7 for 3-5 days before performing the experiments. For the generation of γδ17 T-LTICs, isolated TCRγδ cells were cultured for 3-4 months, and media and IL-7 were refreshed once a week until spontaneous growth was detected in the cultures. γδ17 T-LTICs have been maintained in IL-7 for over 3 years without substantial changes in phenotype in terms of γδ17 T cell markers and cytokine production profile. γδT17-LTICs have resisted at least 3 freeze-thaw cycles. For IL-23/inhibitor treatments, γδT17-LTICs were washed twice in IMDM-10%-2ME to remove IL-7 and cultured for 18h in IMDM-10%-2ME without IL-7. Next day, cells were stimulated with IL-23 (10ng/ml), in presence or absence of inhibitory drugs for 24h. Drugs used in this study: ruxolitinib (0.5μM, Selleckem), Cyclosporin A (Csa, 100nM, Calbiochem), Dasatinib (Selleckem and Med. Chem Express), Bosutinib (Selleckem) and rapamycin (20nM, Tocris).

### Drug repurposing screening

γδT17-LTICs were seeded in 96-well plates (1×10^6^ cells per well) and stimulated with IL-23 (10ng/ml) in presence of the selected pharmacological compounds in three independent biological replicates. The 88 drugs from Plate 1 of the L1300-Selleck-FDA-Approved-Drug-Library were added at 1μM. Control conditions (triplicates) included IL-23+ 0.5% DMSO, IL-23 + Ruxolininib 0.5μM (Selleckem) or DMSO alone. Cells were incubated for 20h before harvesting the supernatant and cells. The amount of IL-17A in the supernatant was assessed by ELISA (ELISA MAX™ Deluxe Set Mouse IL-17A, Biolegend). Cell viability was assessed by staining with Ghost Dye™ Red 780 (Cytek) and flow cytometry analysis. Heatmap representing the IL-17A production was created using Morpheus (https://software.broadinstitute.org/morpheus/).

### In vitro generation of γδ17T

Lymph nodes (LNs) from 10 to 16-week-old C57BL/6 mice were harvested and mechanically disaggregated. Cells were stimulated with αTCRγδ (clone GL3, 50ng/ml, BD), IL-23 (10ng/ml, Miltenyi) and IL-1β (0.1ng/ml, Peprotec) for 5 days in presence/absence of Dasatinib. Cells were washed and cultured for additional 20h in IMDM-10% with IL-7 5ng/ml. Next day, frequencies and numbers of γδ17T cells were determined by flow cytometry.

### In vitro generation of Th17 cells

Lymph nodes (LNs) from 10 to 16-week-old C57BL/6 mice were harvested and mechanically disaggregated. Naïve CD4+ T cells were isolated by negative magnetic sorting. Briefly, cells were incubated with a mixture of biotinylated antibodies (αMHC-II I-A/I-E, clone2G9; αIgM, clone R6-60.2; αCD11c, clone HL3; αCD49b, clone DX5; αTCRγδ, clone GL3 from BD Pharmigen; αF4/80, clone BM8.1; αB220, clone RA3-6B2; αCD19, clone 1D3; αCD11b, clone M1/70; αTER-119, clone TER-119; αCD8a, clone 53-6.7 from Tonbo) diluted in PBS + 0.5%FBS + 2mM EDTA for 30min/ice. Biotinylated αCD25 (clone PC61; Tonbo) and αCD44 (clone IM7; BD Pharmingen) were added during the last 5min for depletion of regulatory and memory T cells. Cells were washed and incubated with streptavidin microbeads (Miltenyi Biotec) for 20min/ice. Cells were washed and passed through MS columns (Miltenyi Biotec) following manufactureŕs instructions. 85% to 95% purity of naïve CD4 cells was routinely obtained. Isolated CD4 T cells were stimulated with plate-bound anti-CD3 (2C11; 5 µg/mL; Tonbo) and anti-CD28 (37.51; 2 µg/mL; eBioscience) in IMDM-10%-2ME supplemented with Th17 polarizing cytokines: TGFβ (3 ng/mL; Prepotec), IL-6 (50 ng/mL; Prepotec), IL-1β (10 ng/mL; Prepotec), IL-21 (10 ng/mL; Prepotec), IL-23 (10 ng/mL; Miltenyi Biotec) for 4-5 days, in presence or absence of different concentrations of Dasatinib. Afterwards, cells were stimulated for 4h with PDBu/Io in presence of Golgi-Plug before assessing IL-17 production by flow cytometry.

### Flow cytometry

Cells were washed twice in PBS and incubated with Ghost Dye-Red780 (Cytek) plus with Fc block (2.4G2, ref. 553142; BD Biosciences) for 30min/ice prior to antibody surface staining for dead cell exclusion. Samples were acquired on a FACSCanto II flow cytometer with DIVA software and on a Aurora 4L with Spectral Flow software (Cytek), and analysed with FlowJo software (Tree Star). Cells were gated according to their forward scatter and side scatter profile, and dead cells excluded based on their staining with the viability dye.

#### Cell surface staining

Cells were washed once in staining solution (PBS, 0.5% FBS, 0.5% bovine serum albumin, and 0.1% sodium azide) and incubated for 20min/ice using manufactureŕs suggested antibody dilutions in staining solution, washed and analysed. γδ17 T cells were electronically gated as CD3^+^TCRγδ^+^CD44^hi^, and CD3^+^TCRγδ^+^CD44^hi^ IL-23R/GFP^+^ when *Il23r*^+/gfp^ reporter mice were used.

#### Cytokine production intracellular staining

After cell stimulation (20h in most cases for IL-23 stimulation), cells were incubated with GolgiPlug (1/1000 dilution, BD Biosciences) for 4h. When indicated, cells were stimulated with phorbol 12,13-dibutyrate (PDBu; 20 ng/ml; Calbiochem) and Ionomycin (Io, 0.5ng/ml, Merck) for 4h in presence of GolgiPlug, or Golgi-Plug alone. Cells were harvested, cell surface stained and fixed for 20 min/RT (IC Fixation buffer eBioscience, ThermoFisher), and incubated for 30min/RT with manufactureŕs suggested antibody dilutions in eBioscience Permeabilization buffer (ThermoFisher). Cells were washed twice in permeabilization buffer and analysed.

#### Phosphoflow

For intracellular staining with anti-phosphospecific antibodies, after stimulations and cell surface staining cells were fixed 30 min/ice followed by 20min/RT, washed and permeabilized in cold 90% methanol (−20°C) for 30min/ice. Cells were then washed in staining solution and incubated with anti-pSTAT3-Y705, pS6-S235/236 or pSrc-Y416 (Cell Signaling Tech) in staining solution for 30 min/RT, washed twice and stained with anti-rabbit-Alexa 647 antibody (Jackson Immunoresearch) for 30 min/RT, washed twice and analysed. For quantification, the mean of fluorescent intensity (MFI) was normalized to the untreated condition to compare different experiments.

#### Cell numbers

Cell numbers were determined by flow cytometry using Accucheck counting beads (ThermoFisher), or using the flow cytometry Aurora 4L counts.

### Imiquimod treatments and analysis

For screening validation, C57BL/6 mice were treated with a cream containing 5% Imiquimod on ears and shaved backs, daily, for 5 days (50mg/day for shaved backs and 10 mg/day for ears; Aldara; Meda Pharma). At the experimental endpoint, skin draining LN (cervical, axillary, brachial and inguinal) were harvested, pooled and mechanically disaggregated. Total lymph node cell suspension were stimulated with IL-23 (10ng/ml) and IL-1β (0.1ng/ml) in presence of different concentrations of Dasatinib and Bosutinib. Cells were incubated for 20h before harvesting the supernatant and cells. The amount of IL-17A in the supernatant was assessed by ELISA (ELISA MAX™ Deluxe Set Mouse IL-17A, Biolegend). For evaluation of Dasatinib therapeutic potential, C57BL/6 and *Il23r*-gfp mice were treated with 5% Imiquimod cream only in the ears, daily, for 7 days (10 mg/day). Dasatinib (Selleckem and MedChemExpress) or control vehicle was administered intraperitoneally (50mg/kg or 10mg/kg) following manufactureŕs instructions (prepared in DMSO and diluted in PBS 30%PEG300, 5%Tween80 from Selleckem). For topical administration, a 150mg/ml Dasatinib stock in DMSO was used to prepare a 1% Dasatinib solution (w/v) in PBS with 30% PEG300 and 5% Tween80. A 20ul drop of this solution was spread on the ear and air-dried for 30s, three times. IMQ dose was applied 30-60min later. At the experimental endpoint, left ear was used for immunohistochemistry analyses, right ear and skin draining LN were used for flow cytometry analysis. Skin draining LN (cervical, axillary and brachial) were harvested, pooled and mechanically disaggregated for flow cytometry analysis. Right ears were split in two halves, cut into pieces and digested for 45min/37°C in RPMI containing Liberase TM (83 mg/ml; Roche), DNase I (100 mg/ml; Roche) and Collagenase IV (0.5 mg/ml; Sigma). Undigested skin pieces were further subjected to tissue disruption using 7 mm stainless steel beads (Qiagen) and a TissueLyser LT (20 oscilations/5 min; Qiagen), and processed for flow cytometry analysis. Left ears were rapidly immersed and fixed in 4% paraformaldehyde and submitted to the Histology Facility at CNB-CSIC for the histological preparation of biological samples. Ears were embedded in paraffin, and skin slices (4-5 mm thick taken 200mm apart) were stained with hematoxylin and eosin (H&E). For IHC staining, skin sections were deparaffinized, boiled in antigen retrieval solution (10mM sodium citrate, 0,05% Tween 20, pH6). Slides were developed with DAB substrate (Dako K3468) and then counterstained with Mayer’s Hematoxylin. Images were captured using an Axio Imager. M1 vertical, 5x objective, with a Leica DMC6200 CMOS Color camera. The thickness of the epidermal layer was measured at multiple sections and sites, randomly chosen in a blind manner (8 sections per mouse, 20 measures per section), using ImageJ software.

### Western blotting

Tγδ17-LTICs cells were lysed at 20-30×10^6^/ml in RIPA buffer (100 mM HEPES, pH 7.4, 150 mM NaCl, 1% NP40, 0.1% SDS, 0.5% sodium deoxycholate, 10% glycerol, 1 mM EDTA, 1 mM EGTA, 1 mM TCEP (Pierce), and protease and phosphatase inhibitors (Roche). Protein samples were mixed with NuPAGE LDS sample buffer (Life Technologies) supplemented with TCEP as reducing agent (25 mM; Sigma-Aldrich). Protein lysates were separated by SDS-PAGE polyacrylamide gel electrophoresis and then transferred to nitrocellulose membrane (Amersham) using standard conditions (Mini-PROTEAN tetra cell system; Bio-Rad). Membranes were blocked with 5% (w/v) non-fat dried skimmed milk powder in PBS containing 0.05% Tween 20. Membranes were probed with the following primary antibodies: pSTAT3-Y705, PanSTAT3, pSrc-Y416, pS6K1-T389, pS6-S235/236, PanS6, pAkt-S473 (all purchased from Cell Signaling) and SMC1 (Bethyl). Primary antibodies were incubated for 20h at 4°C (diluted in PBS 0.05% Tween20 5%BSA) and detected using peroxidase-conjugated secondary antibodies (Goat anti-Rabbit-Ig-HPO and Goat anti-Mouse-Ig-HPO, Cell Signaling), and chemiluminescence detected with the ImageQuant LAS-4000 imaging system (Fujifilm).

### IGP dataset

This work benefitted from data assembled by the ImmGen consortium (Heng, Painter, and Immunological Genome Project Consortium, 2008). Dataset: Tgd.g2+d17.LN (sorted on TCRvg2+TCRd+CD24-CD27-CD4-CD8-) by Kang Lab. University of Massachusetts Medical School, Dataset GSE109125_Normalized_Gene_count.

### Lentiviral production of shBlk

Predesigned shRNA against Blk cloned into pLKO.1-TRC1 lentiviral vector (NM_007549/ TRCN0000023410; Sigma MISSION shRNA Library) was subcloned into the LV-GFP lentiviral vector as a SphI/SacII fragment. LV-GFP vector was a gift from Elaine Fuchs (Addgene plasmid #25999). Restriction enzymes were obtained from New England Biolabs. For lentiviral particle production, LV-GFP and LV-GFP-shBlk were transiently transfected in HEK293 cells together with the packaging vector psPAX2 and the envelope vector pMD2.G (gifts from Didier Trono; Addgene plasmids #12260 and #12259, respectively) using Polyethylenimine (PEI; Sigma). Next day, culture supernatants were replaced with fresh media. 48h and 72h after transfection, culture supernatants were harvested and concentrated by ultracentrifugation (26.000 rpms/2 h/4 C°). Concentrated lentiviral particles were kept at −80 C° until use. γδ17 T-LTICs cells were stimulated for 20 h with IL-23 prior to transduction with lentiviral particles in presence of Polybrene (10 µg/mL). Culture plates were spun at 650g/1h/32°C. Next day, cells were washed and resuspended in fresh medium containing IL-7. 7-10 days later, cells were stimulated with IL-23 and transduction efficiency, pS6 and IL-17 production were determined by flow cytometry. HEK293 cells were maintained in DMEM (Invitrogen) supplemented with L-glutamine, 10% (v/v) FBS, penicillin (50 U/mL) and streptomycin (50 mg/mL).

### Statistical Analysis

All datasets were subjected to D’Agostino & Pearson omnibus normality test to determine Gaussian distribution. The datasets did not pass normality test and accordingly, the statistical significances were obtained using the non-parametric MannWhitney two-tailed t-test and Kruskal-Wallis test with Dunńs correction for multiple comparisons. Graphad Prism v.6 and v.8 were used for statistical analysis.

## Results

### Development a γδ17 T Cell-Based System to Study IL-23-Driven Type 3 Immunity

To screen pharmacological compounds that interfere with IL-23-induced IL-17 production, we developed a robust cellular model of type 3 immunity based on γδ17 T cells. These cells were expanded in long-term cultures in the presence of IL-7 (γδ17 T-LTICs, Long Term IL-7 Cultures) (Michel *et al*., 2012), using γδ17 T cells isolated from *Il23r*^wt/gfp^ reporter mice (Awasthi *et al*., 2009). After 3–4 months, the cells began to proliferate spontaneously and could be maintained in culture for more than three years. Long-term in vitro cultures often lead to adaptation and phenotypic drift from primary features. To assess this possibility, we compared γδ17 T-LTICs with *ex vivo* γδ17 T cells and found that the cultured cells retained the expression of key phenotypic markers: high CD44, absence of CD27, and expression of RORγt and IL-23R (**Fig. 1a**). Functionally, γδ17 T-LTICs preserved the ability to produce IL-17A, IL-22, IL-17f, and GM-CSF (**Supplemental Fig. 1a**). Upon IL-23 stimulation, they upregulated IL-17A, IL-22, and GM-CSF production (**Fig. 1b and Supplemental Fig. 1a**) and induced key events of in IL-23 signalling cascade such as STAT3-Y705 and S6-S235/236 phosphorylation (**Supplemental Fig. 1b**). These cells displayed a low proliferation rate in the presence of IL-7 (**Fig. 1c**). To validate this cell system for drug screening, we tested two psoriasis therapies: the Jak2 inhibitor Ruxolitinib and the calcineurin inhibitor Ciclosporin A (Reid and Griffiths, 2020). Both drugs effectively suppressed IL-23-induced IL-17A production (**Fig. 1d**), demonstrating that γδ17 T-LTICs represent a reliable model for identifying compounds that modulate IL-23-driven IL-17A production. Thus, γδ17 T-LTICs constitute a stable and functionally relevant *in vitro* model of type 3 immunity, and they can be a valuable platform for screening pharmacological inhibitors of the IL-23/IL-17 axis.

**Figure 1.**
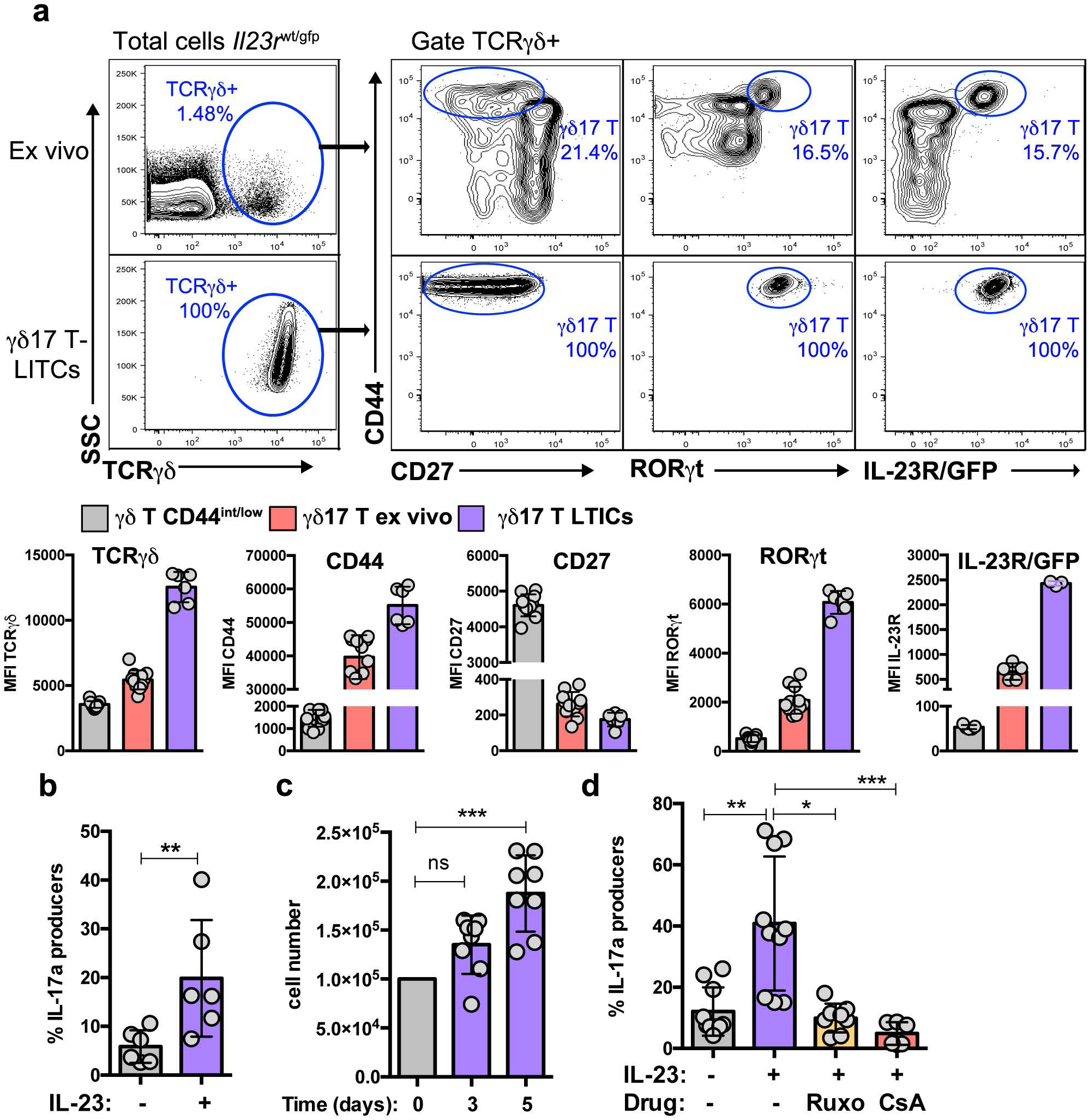
γδ17 T Long-Term IL-7 Cultures (γδ17 T-LTICs) maintain γδ17 T cell markers and IL-23 responsiveness. γδ T cells isolated from the lymph nodes of *Il23r*^wt/gfp^ reporter mice were cultured in presence of IL-7. 4-12 months later, cells were analysed for expression of γδ17 T markers and IL-23 response. **a)** Expression of γδ17 T cell markers in ex vivo lymph node γδ T cells and γδ17 T-LTICs. Cells were stained with the indicated antibodies and analysed by flow cytometry. Cells were stained with the indicated antibodies and analysed by flow cytometry. Left dot plots show TCRγδ expression. Right dot plots show expression of CD44, CD27, RORγt, and IL-23R/GFP in gated TCRγδ⁺ cells. Graphs show the mean fluorescence intensity (MFI) of these markers (mean±sd, n = 5 different animals for ex vivo γδ T, n = 3 independent γδ17 T-LTICs cultures). **b)** γδ17 T-LTICs were stimulated with IL-23 for 20h before assessing IL-17 production by flow cytometry. Graph shows the frequency of IL-17A⁺ cells (mean±sd, n = 6 independent experiments). **c)** γδ17 T-LTICs were cultured in presence of IL-7, and cell number was assessed at day 3 and 5. Graph shows cell numbers normalized to day 0 (mean±sd, n = 8 independent experiments). **d)** γδ17 T-LTICs were stimulated with IL-23 for 20h in presence of the Jak2 inhibitor Ruxolitinib (Ruxo) or the calcineurin inhibitor ciclosporin A (Csa), before assessing IL-17 production by flow cytometry. Graph shows the frequency of IL-17A⁺ cells (mean±sd, n = 10 independent experiments). Statistical analysis. b) Mann-Whitney test, **p=0.0087, c, d) Kruskal-Wallis test with Dunńs correction for multiple comparisons. c)***p=0.0003, ns=no significant d)**p=0.0068, *p=0.0159,***p=0.0001.

### Drug-repurposing screening identifies Src/Abl inhibitors Dasatinib and Bosutinib as suppressors of IL-23 signalling

We stimulated γδ17 T-LTICs with IL-23 in 96-well plates, in presence of 88 FDA-approved drugs for 20h in three independent biological replicates (one drug per well). Next day we determined IL-17A production and cell viability. As control conditions we used unstimulated cells, IL-23 alone and IL-23+Ruxolitinib, a Jak2 inhibitor that block IL-23 signalling (Parham *et al*., 2002). The plate layout, compound list and raw screening results can be found in **Supplemental Table I**, and **Fig. 2a** shows the normalized results as a heatmap. The reference inhibitor Ruxolitinib reduced IL-17A production by 70%. Of the 88 tested compounds, 4 showed a high cellular toxicity (viability below 60% of control condition), and 10 showed a strong and consistent reduction of IL-23-induced IL-17 production (50% reduction) (**Fig. 2b)**. Most of the identified compounds are approved for cancer treatment, and among them we found rapamycin, a mTORC1 inhibitor previously linked to IL-23 signalling (Cai *et al*., 2019; Cibrian *et al*., 2020). We also found drugs targeting VEGFR/PDGFR (Vandetanib, Sunitinib and Pazopanib), Src/Abl kinases (Dasatinib and Bosutinib), c-Met kinase (Cabozantinib and Crizotinib), estrogen/progesterone receptors (Dienogest) and HDACs (Vorinostat). Of the screening hits, we selected the Src/Abl inhibitors Dasatinib and Bosutinib for further analysis, as these kinases are well-known regulators of T cell signalling that have not been previously linked to IL-23 signalling (Hwang *et al*., 2020). We performed a dose response curve for IL-17 production in γδ17 T-LTICs, confirming the strong inhibition of IL-17A production (**Fig. 2c,d**). Next, we wanted to examine if Dasatinib and Bosutinib were capable of inhibiting IL-17 production in pathogenic cells obtained from the psoriasis-like model induced by Imiquimod (IMQ) treatment that is mediate by the IL-23/IL-17 axis (van der Fits *et al*., 2009). In this pro-inflammatory context, IL-1β synergizes with IL-23 to induce the production of large amounts of IL-17 and other mediators in autoimmune diseases (Sutton *et al*., 2009; Coccia *et al*., 2012; Cai *et al*., 2014). We stimulated lymph node cells obtained from IMQ-treated animals with the combination of IL-23 and IL-1β in presence of Dasatinib or Bosutinib, and determined that Dasatinib but not Bosutinib inhibited IL-17 production in cells obtained from IMQ model (**Fig. 2e**). We also assessed the impact of Dasatinib in the generation and expansion of IL-17-producing cells *in vitro*. γδ T cells commit to IL-17 production during thymic development, and in response to immune challenges, extrathymic signals such as TCR stimulation, IL-23 and IL-1β induce the differentiation of naïve γδ T cells into γδ17 effector cells (Muschaweckh, Petermann and Korn, 2017; Papotto *et al*., 2017). We stimulated lymph node cells with anti-TCRγδ, IL-23, and IL-1β in presence of Dasatinib for 5 days, and observed a marked decrease in the number of *in vitro* generated γδ17 cells (**Fig. 2f**). We also performed in vitro differentiation assays of naïve CD4 T cells under Th17-polarizing conditions, observing a marked reduction in Th17 differentiation in the presence of Dasatinib (**Fig. 2g**). Thus, Dasatinib impaired the generation of IL-17-producing γδ T and CD4 T cells. To summarise, the drug-repurposing screening using the γδ17 T-LTICs identified FDA-approved compounds that reduce IL-23-induced IL-17A production, including the Src/Abl kinase inhibitors Dasatinib and Bosutinib. These compounds, primarily approved for cancer treatment, demonstrated potent inhibitory effects in vitro assays, with Dasatinib showing greater efficacy.

**Figure 2.**
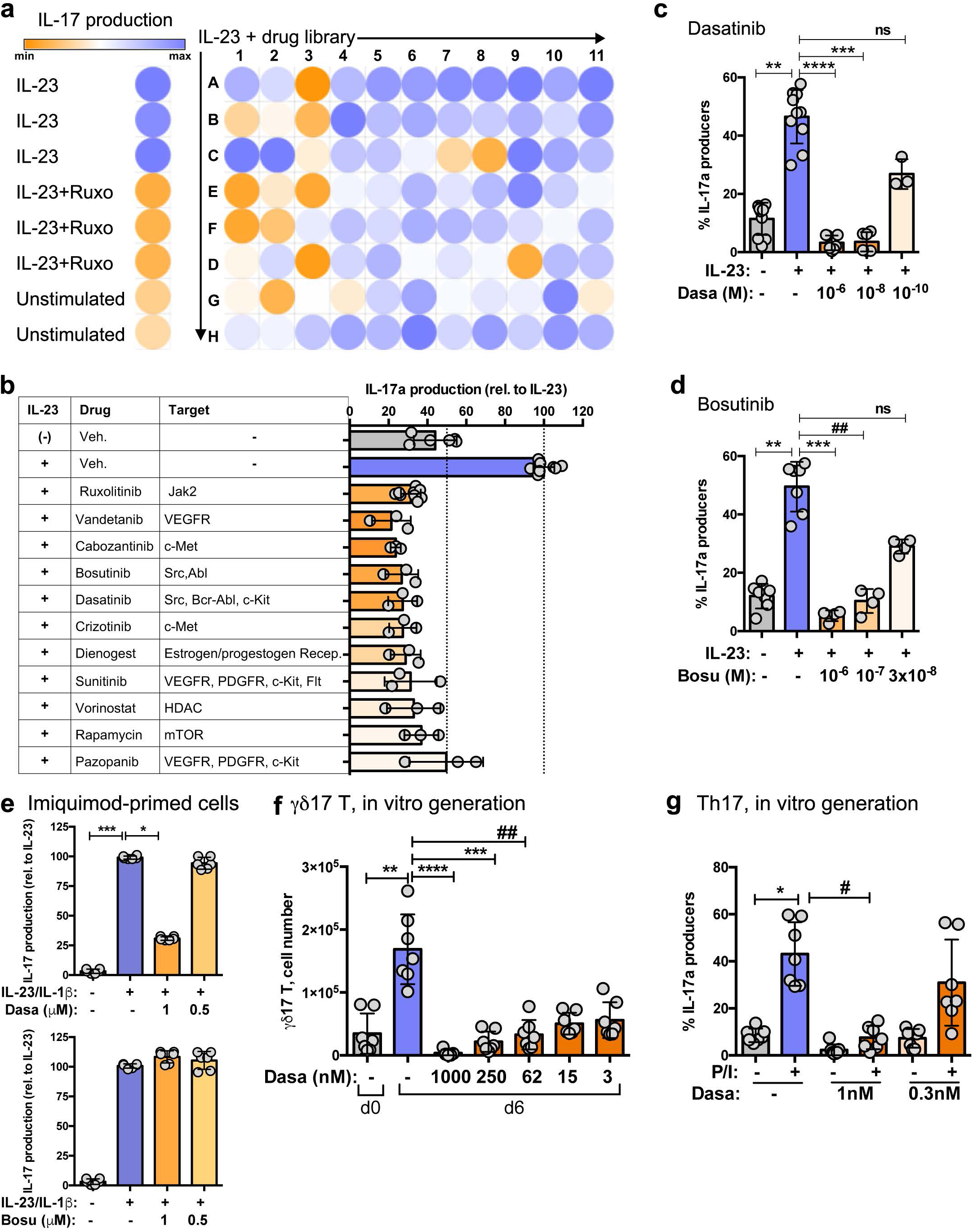
Drug-repurposing screening identifies dual Src/Abl inhibitors as suppressors of IL-23 response. γδ17 T-LTICs were seeded in 96-well plates and stimulated with IL-23 in the presence of FDA-approved drugs (one compound per well, 1 μM) or DMSO for 20 h. The Jak2 inhibitor Ruxolitinib (Ruxo) was added as a reference inhibitor. IL-17 production was measured by ELISA, and cell viability was assessed by flow cytometry. **a)** Heatmap shows IL-17 production in each well, normalized to the IL-23 condition (averaged value of n = 3 screening replicates). **b)** List of top 10 suppressors of IL-23-induced IL-17 production. Table shows drugs name and described targets, and graphs show the normalized IL-17 production (mean±sd, n= 3 screening replicates). **c, d)** γδ17 T-LTICs were stimulated with IL-23 for 20h in presence of the indicated concentrations of the Src/Abl inhibitors Dasatinib (c) or Bosutinib (d), before assessing IL-17 production by flow cytometry. Graphs show the frequency of IL-17A⁺ cells (mean±sd, n=3-10). **e)** WT mice were treated with Imiquimod (IMQ) for 5 days. Skin-draining lymph node cell suspensions were stimulated with IL-23 and IL-1β for 20h in presence of the indicated concentrations of Dasatinib and Bosutinib. IL-17 production was assessed next day by ELISA. Graphs show IL-17 production, normalized to the IL-23 condition (mean±sd, n= 6 animals). **f)** Total lymph node cells were stimulated with αTCRγδ/IL-23/IL-1β for 5 days in presence of the indicated concentrations of Dasatinib. Graph shows the number of γδ17 T cells (gated as CD3+TCRγδ+CD44hi) assessed by flow cytometry (as mean±sd, n=7 different mice). **g)** Isolated naïve CD4 T cells were stimulated in plate-coated αCD3/αCD28 and Th17 polarising cytokines (IL-6, TGFβ, IL-1β, IL-21 and IL-23), in presence of the indicated concentrations of Dasatinib. After 4 days, cells were stimulated with PDBu/Io in presence of Golgi-Plug for 4h before assessing IL-17 production by flow cytometry. Graph shows the frequency of IL-17-producing cells (as mean±sd, n=6-7 mice). Statistical analysis was assessed using Kruskal-Wallis test with Dunńs correction for multiple comparisons. c). **p=0,0073, ****p<0.0001, ***p=0.0002, ns=no significant. d) **p=0.0063, ***p=0.0003, ##p=0.0092, ns=no significant. e) ***p=0.0002, *p=0.0338. f)**p=0.0052, **** p<0.0001, ***p= 0.0009, ##p= 0.0090, g)*p= 0.0266, #p= 0.0119.

### Intraperitoneal Dasatinib administration ameliorates skin inflammation in the psoriasis-like Imiquimod model

The marked reduction of IL-17 production by Dasatinib in vitro prompted us to examine its therapeutic potential in the IMQ model of skin inflammation. We administered daily intraperitoneal injections of Dasatinib (50 mg/kg; 7 days) and analysed different inflammation parameters. We assessed keratinocyte hyperproliferation by measuring epidermal thickness, and detected over 50% reduction in IMQ/Dasatinib-treated animals compared with IMQ alone (**Fig. 3a**). Flow cytometry analysis of skin leukocyte infiltrate (gating strategy shown in **Supplemental Fig. 2**) revealed a significant reduction in neutrophils (Ly6G^+^) and inflammatory monocyte-macrophages (CD64^+^Ly6C^+^) (**Fig. 3b**). We also observed a pronounced reduction in TCRγδ cells (**Fig. 3c**). Within the TCRγδ population, two subpopulations -dermal (TCRγδ^int^) and epidermal (TCRγδ^hi^)- can be distinguished, with the dermal subset being the primary IL-17 producers in this skin inflammation model (Cai *et al*., 2011, 2014). Dasatinib treatment reduced both subpopulations, as well as the recruitment of CD4 T cells (**Fig. 3c**). In the IMQ model, Th17 and γδ17 T cells expand in the draining lymph nodes and spleen before migrating to inflamed skin, where they contribute to lesion development (van der Fits *et al*., 2009; Gray *et al*., 2013; Cai *et al*., 2014; Ramírez-Valle, Gray and Cyster, 2015). Dasatinib-treated mice showed a significant decrease in total TCRγδ cells and in the TCRγδ^+^CD44^hi^ subpopulation, which comprises the γδ17 T cells (Schmolka *et al*., 2013) (**Fig. 3d**). CD4 T cell numbers in the draining lymph nodes were also reduced (**Fig. 3d**). Finally, in vitro stimulation with phorbol esters and ionomycin confirmed a significant decrease in IL-17-producing γδ and CD4 T cells in the skin-draining lymph nodes of Dasatinib-treated mice compared with vehicle controls (**Fig. 3e,f**). Overall, Dasatinib significantly reduced leukocyte infiltration, epidermal hyperproliferation, and IL-17 production in the IMQ model of skin inflammation.

**Figure 3.**
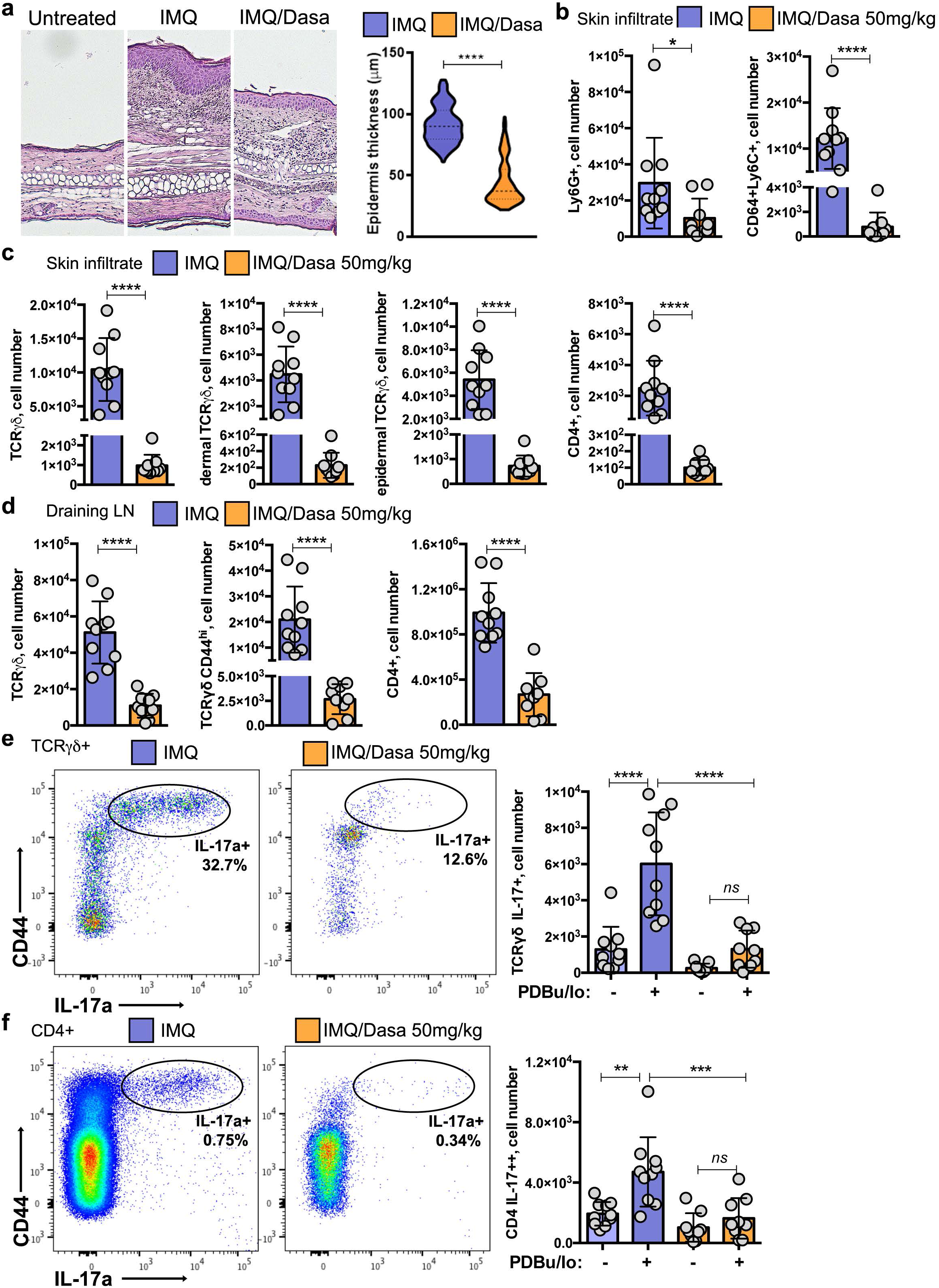
Dasatinib reduces epidermal thickening, skin infiltrate and T cell expansion in IMQ skin inflammation model. C57BL6/J WT animals were treated with Imiquimod (IMQ) for 7 days on ears, together with a daily intraperitoneal injection of Dasatinib (50mg/kg) or control vehicle. On day 7, ear skin and draining lymph nodes were processed for histological and flow cytometry analysis. All graphs represent pooled data from 2 independent experiments (as mean±sd, n = 9-10 animals per treatment). **a)** Representative sections of H/E staining of untreated, IMQ and IMQ/Dasa-treated ears. Violin plot shows the epidermal layer thickness. **b)** Graphs show the number of neutrophils (Ly6G^+^) and inflammatory monocyte-macrophages (Ly6G^neg^CD64^+^Ly6C^+^) in the skin infiltrate. **c)** Graphs show the number of TCRγδ (CD3^+^TCRγδ^+^), dermal TCRγδ (TCRγδ^int^), epidermal (TCRγδ^hi^) and CD4 (CD3^+^TCRγδ^neg^CD4^+^) T cells in the skin infiltrate. **d)** Graphs show the number of TCRγδ (CD3^+^TCRγδ^+^), γδ17 T (CD3^+^TCRγδ^+^CD44^hi^) and CD4 cells (gated as CD3^+^TCRγδ^neg^CD4^+^) in skin draining lymph nodes. **e, f)** Skin draining lymph node cells were stimulated with PDBu/Io of left unstimulated for 4h in presence of Golgi-Plug before assessing IL-17A production by flow cytometry. **e)** Representative dot plots show CD44 and IL-17A production upon PDBu/Io stimulation in TCRγδ cells. Graph shows the number of TCRγδ^+^IL-17^+^ cells in the indicated conditions. **f)** Representative dot plots show CD44 and IL-17A production upon PDBu/Io stimulation in CD4 T cells. Graph shows the number of CD4^+^IL-17^+^ cells in the indicated conditions. Statistical analysis was assessed using Mann-Whitney t-test (a-d) and Kruskal-Wallis test with Dunńs correction for multiple comparisons (e-f). a) **** p< 0.0001. b) *p= 0.0133, ****p<0.0001. c) ****p< 0.0001. d) ****p< 0.0001. e) ****p< 0.0001, ns, no significant. f) **p=0.0010, ***p=0.0004, ns, no significant.

Next, we investigated the impact of a fivefold lower dose of Dasatinib (10 mg/kg) on IMQ-induced skin inflammation. To track type 3 cells (mainly γδ17 and Th17 T cells) and avoid the cell death induced by PDBu/Io stimulation, we used the *Il23r*^wt/GFP^ reporter mouse, which serves as a surrogate reporter for RORγt expression and thus identifies cells capable of producing IL-17 (Awasthi *et al*., 2009; Yang *et al*., 2014). In these experiments, we did not observe a consistent reduction in epidermal thickness (**Fig. 4a**). We noted a decrease in myeloid infiltration that did not reach statistical significance (**Fig. 4b**). Regarding lymphoid cell infiltrate, the 10 mg/kg dose markedly reduced the number of dermal γδ T cells, particularly the IL-23R-expressing γδ T cells, as well as CD4 T cells (**Fig. 4c**). Analysis of the skin-draining lymph nodes revealed a strong reduction in γδ T cells, specifically in the IL-23R-expressing subpopulation (**Fig. 4d**). The numbers of CD4 T cells and in particular the IL-23R^+^ CD4 T cells were also reduced (**Fig. 4e**). In summary, Dasatinib administration primarily reduced lymphoid cell infiltration in the skin and decreased the generation and expansion of IL-17-producing CD4 and γδ17 T cells in the secondary lymphoid organs of IMQ-treated animals.

**Figure 4.**
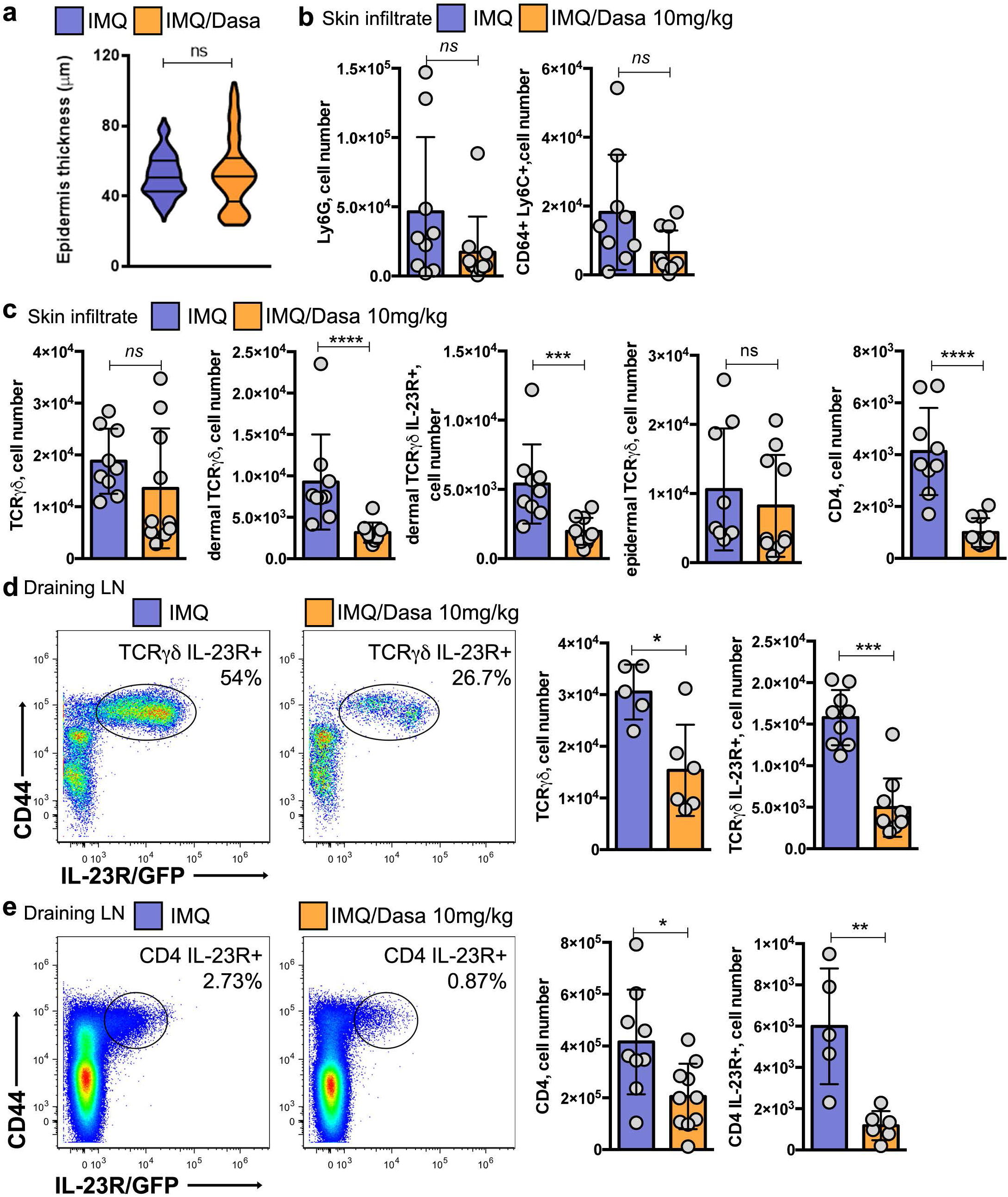
Low-dose Dasatinib attenuates infiltration and expansion of IL-17-producing T cells in IMQ-Model in IMQ skin inflammation model. *Il23r*^wt/gfp^ reporter animals were treated with Imiquimod (IMQ) for 7 days on ears, together with a daily intraperitoneal injection of Dasatinib (10mg/kg) or control vehicle. On day 7, ear skin and draining lymph nodes were processed for histological analysis and flow cytometry. All graphs represent pooled data from 2 independent experiments (as mean±sd, n=9-10 animals per treatment). **a)** Violin plot shows the epidermal layer thickness in IMQ and IMQ/Dasa-treated ears. **b)** Graphs show the number of neutrophils (Ly6G^+^) and inflammatory monocyte-macrophages (Ly6G^neg^CD64^+^Ly6C^+^) in the skin infiltrate. **c)** Graphs show the number of TCRγδ (CD3^+^TCRγδ^+^), dermal TCRγδ (TCRγδ^int^), dermal TCRγδ IL-23R/GFP^+^, epidermal (TCRγδ^hi^) and CD4 T cells (CD3^+^TCRγδ^neg^CD4^+^) in the skin infiltrate. **d)** Representative dot plots show CD44 and IL-23R/GFP expression in γδ T cells in the skin draining lymph nodes. Graphs show the number of total TCRγδ cells (CD3^+^TCRγδ^+^) and γδ17 T cells (CD3^+^TCRγδ^+^CD44^hi^IL-23R/GFP^+^). **e)** Representative dot plots show CD44 and IL-23R/GFP expression in CD4 T cells in the skin draining lymph nodes. Graphs show the number of total CD4 cells (CD3^+^TCRγδ^neg^CD4^+^) and CD4 IL-23R^+^ cells (CD3^+^CD4^+^CD44^hi^IL-23R/GFP^+^). Statistical analysis was assessed using Mann-Whitney test. ns, no significant. c) ****p<0.0001, ***p=0.0003, d) *p=0.0303, ***p=0.0002, e) *p=0.0133 **p=0.0043.

### Topical Dasatinib administration reduces Imiquimod-skin inflammation

One of the advantages of pharmacological compounds is their potential for oral or topical administration, which represents a great improvement on patients’ quality of life. To explore this possibility, we evaluated the therapeutic potential of topical Dasatinib administration. In these experiments, we detected a slight but significant 5% decrease in the engrossment of the epidermal layer induced by IMQ treatment (**Fig. 5a**). We did not observe significant changes in the number of myeloid cells in the inflamed skin (**Fig. 5b**). However, we noted a specific reduction in lymphoid populations (dermal and epidermal γδ T cells, and CD4 T cells) (**Fig 5c**). To determine if topical Dasatinib had only local effects on the skin or if it could have potential systemic effects, we examined the number of γδ and CD4 T cells in the skin draining lymph nodes. These experiments showed a non-significant decrease in total γδ T cells, and a significant decrease in IL-23R-expressing γδ17 T cells (**Fig. 5d**). Regarding the CD4 population, we observed a slight decrease in the total number of CD4 T cells, with no consistent changes in the IL-23-expressing subpopulation (**Fig. 5e**). Thus, topical Dasatinib administration reduced skin infiltration of lymphoid cells and selectively decreased the generation and expansion of IL-17-producing CD4+ and γδ17 T cells in secondary lymphoid organs. These findings underscore Dasatinib’s potential to block type 3 immune responses in vitro and in a psoriasis-like skin inflammation model, supporting its therapeutic potential in inflammatory conditions.

**Figure 5.**
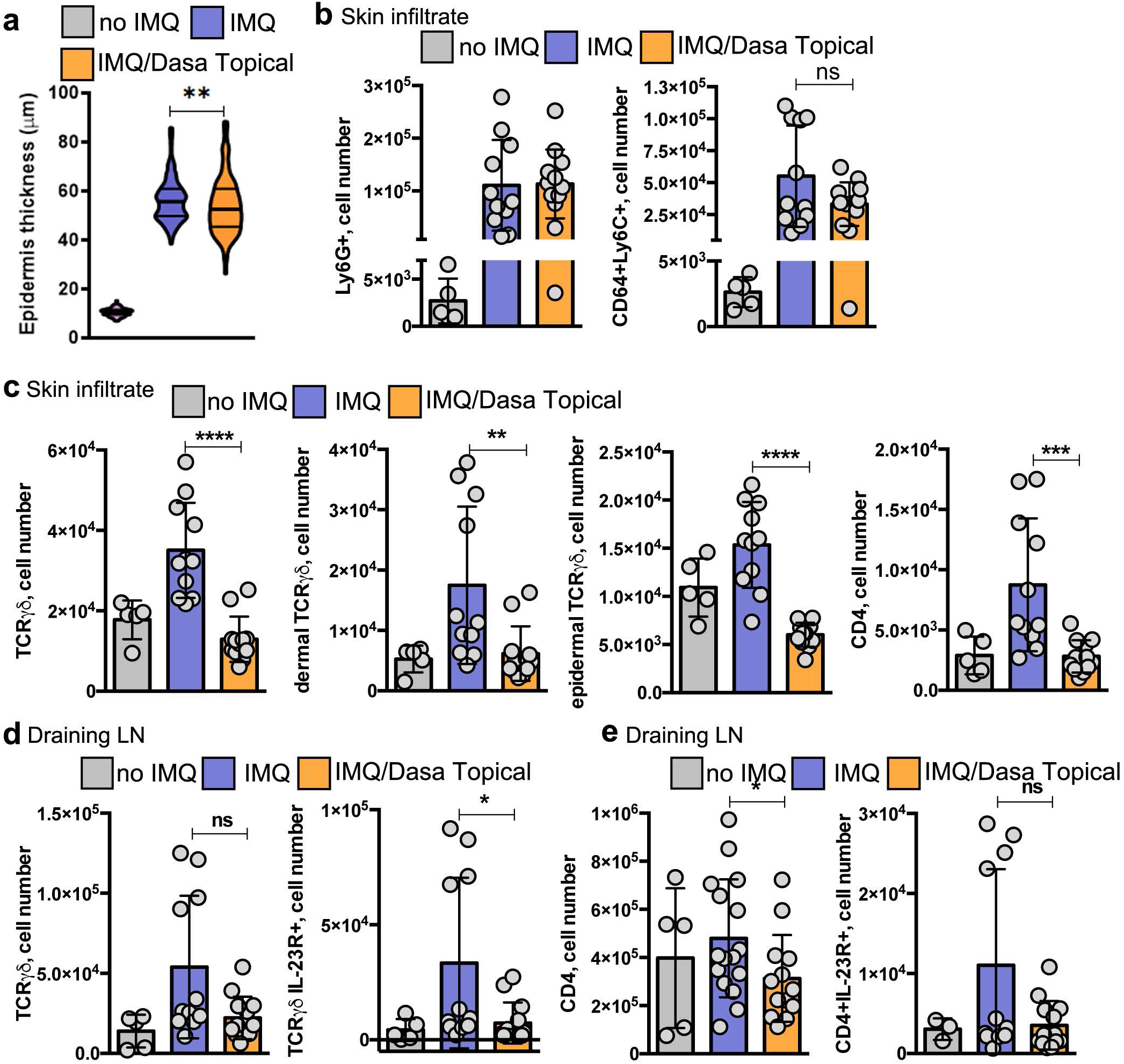
Topical Dasatinib reduces skin lymphoid infiltrate and expansion of IL-17-producing T cells in IMQ-induced psoriasis-like model. *Il23r*^wt/gfp^ reporter animals were treated with Imiquimod (IMQ) for 7 days on ears, together with topical administration of 1% Dasatinib solution (w/v) or control vehicle or left untreated. On day 7, ear skin and draining lymph nodes were processed for histological analysis and flow cytometry. All graphs represent pooled data from 2 independent experiments (as mean±sd, n=11-12 animals per treatment). **a)** Violin plot shows the epidermal layer thickness in untreated (no IMQ), IMQ and IMQ/Dasa-treated ears. **b)** Graphs show the number of neutrophils (Ly6G^+^) and inflammatory monocyte-macrophages (Ly6G^neg^CD64^+^Ly6C^+^) in the skin infiltrate. **c)** Graphs show the number of TCRγδ (CD3^+^TCRγδ^+^), dermal TCRγδ (TCRγδ^int^), epidermal (TCRγδ^hi^) and CD4 (CD3^+^TCRγδ^eg^CD4^+^) T cells in the skin infiltrate. **d)** Graphs show the number of total TCRγδ cells (CD3^+^TCRγδ^+^) and γδ17 T cells (CD3^+^TCRγδ^+^CD44^hi^IL-23R/GFP^+^) in the skin draining lymph nodes. **e)** Graphs show the number of total CD4 cells (CD3^+^TCRγδ^neg^CD4^+^) and CD4 IL-23R+ cells (CD3^+^CD4^+^CD44^hi^IL-23R/GFP^+^) in the skin draining lymph nodes. Statistical analysis was assessed using Mann-Whitney (untreated samples were included for reference but not considered in the statistical analysis). ns, no significant. a)**p=0.0044, c)****p<0.0001, **p=0.0056, ***p=0.0005 d) *p=0.0129, e) *p=0.0459.

### Molecular mechanisms: Dasatinib and Bosutinib block IL-23-mediated mTORC1/mTORC2 activation

To investigate the molecular mechanisms underlying the inhibitory effect of Dasatinib and Bosutinib, we examined their impact on the IL-23 signalling cascade. IL-23 signalling relies on Jak2/Tyk2-mediated phosphorylation of STAT3 (Parham *et al*., 2002) and mTORC1 activation (Cai *et al*., 2019; Cibrian *et al*., 2020). Accordingly, the Jak2 inhibitor Ruxolitinib and the mTORC1 inhibitor Rapamycin effectively blocked IL-23-induced IL-17 production in primary γδ17 T cells (**Fig. 6a**). Moreover, this signalling network is conserved in γδ17 T-LTICs (**Fig. 6b**). To gain insights into the molecular mechanisms, we treated γδ17 T-LTICs with increasing concentrations of Dasatinib or Bosutinib and analysed their impact on the phosphorylation of STAT3-Y705 (Jak2 substrate), and S6K-T389 (mTORC1 direct substrate) and S6-S235/236 (S6K direct substrate and thus, mTORC1indirect substrate). To confirm the efficacy of the Src/Abl inhibitors, we used an anti-phosphospecific antibody against Src phosphorylated at tyrosine 416 (pSrc-Y416), a site within the kinase activation loop whose phosphorylation enhances enzymatic activity. This antibody cross-reacts with various Src family members, serving as a marker of Src family kinase activation. Both Dasatinib and Bosutinib treatment inhibited pSrc-Y416 in a dose-dependent manner, while we did not detect clear changes in pSrc-Y416 in response to IL-23 treatment. Dasatinib and Bosutinib treatments did not affect pSTAT3-Y705, while they strikingly blocked IL-23-induced mTORC1 activation, as evidenced by reduced pS6K-T389 and pS6-S235/236 levels (**Fig. 6c,d**). IL-23 signalling has been also shown to rely on mTORC2 activation (Cai *et al*., 2019). Thus, we explored the impact of Dasatinib and Bosutinib on mTORC2 activation monitoring the phosphorylation of the Akt-S473 residue (pAkt-S473). In these experiments we shown that IL-23 increased pAkt-S473, and the treatment with both inhibitors prevented Akt phosphorylation (**Fig. 6e**). Thus, Dasatinib and Bosutinib inhibited IL-23-dependent mTORC1 and mTORC2 activation without affecting STAT3 phosphorylation.

**Figure 6.**
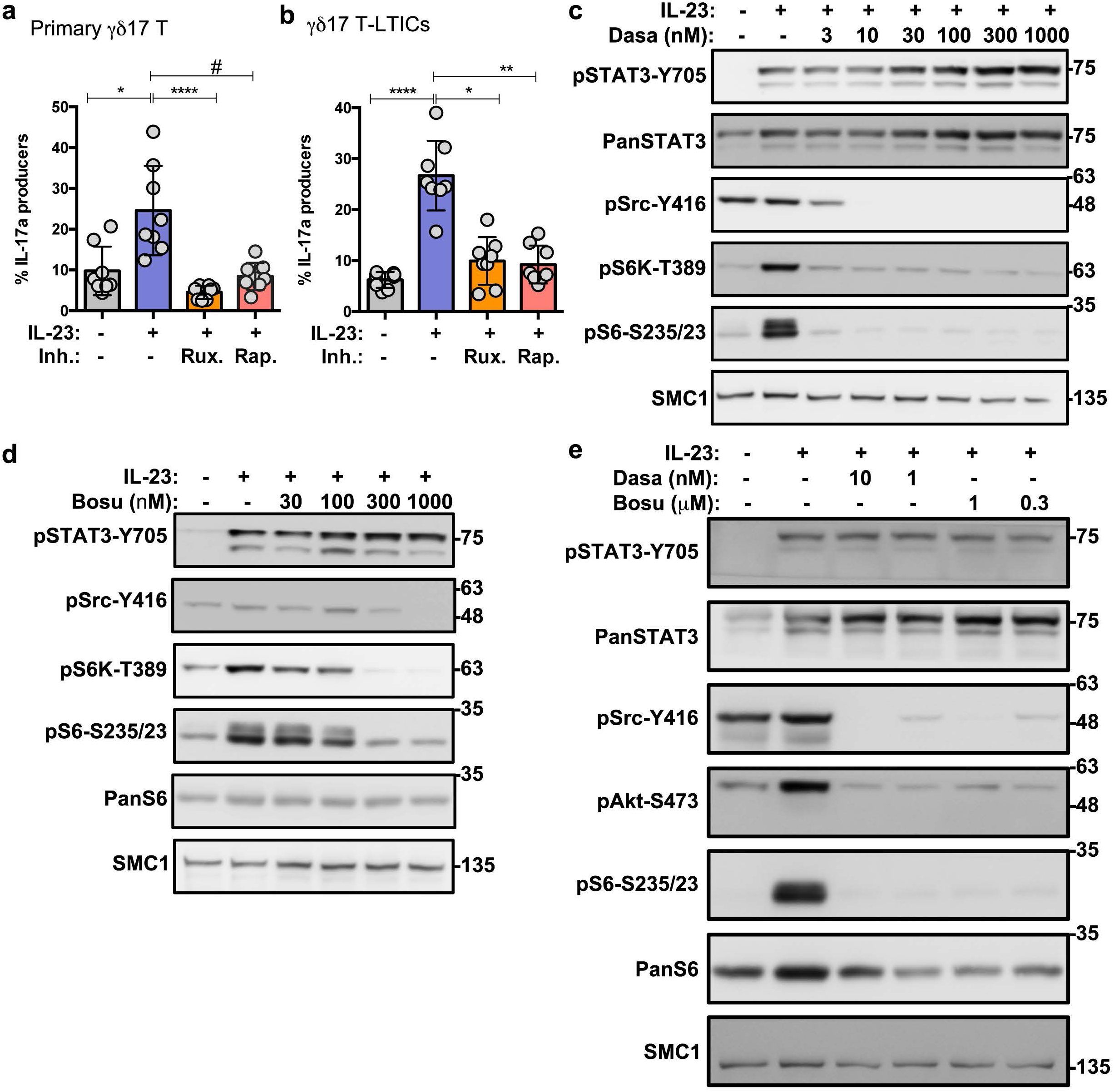
Dasatinib and Bosutinib impair IL-23-mediated activation of mTORC1 and mTORC2. **a**) Primary γδ17 T cells were stimulated with IL-23 for 20h in presence of the Jak2 inhibitor ruxolitinib (Rux.) and the mTORC1 inhibitor rapamycin (Rap.). Graph shows the frequency of IL-17A-producing cells, determined by flow cytometry (mean±sd, n=8). **b)** γδ17T-LTICs were treated and processed as in a). Graph shows the frequency of IL-17A-producing cells, determined by flow cytometry (mean±sd, n=8). **c, d, e)** γδ17T-LTICs were stimulated with IL-23 for 20h in presence of the indicated concentrations of Dasatinib (Dasa.) and Bosutinib (Bosu.), and processed for western-blot analysis. **c, d)** Western-blot analysis of the impact of **c)** Dasatinib and **d)** Bosutinib on: Jak2 substrate pSTAT3-Y705, Src substrate pSrc-Y416, mTORC1 substrate pS6K1-T389, S6K1 substrate pS6-S235/236. PanSTAT3 and SMC1 were used as loading controls. Representative of n= 3 independent experiments. **e)** Western-blot analysis of the impact of Dasatinib and Bosutinib on: Jak2 substrate pSTAT3-Y705, Src substrate pSrc-Y416, mTORC2 substrate pAkt-S473, S6K1 substrate pS6-S235/236. PanSTAT3, PanS6 and SMC1 were used as loading controls. Representative of n=4 independent experiments. Statistical analysis was assessed using Kruskal-Wallis test with Dunńs correction for multiple comparisons a) *p=0.0354, ****p< 0.0001, #p=0.0477, b)****p< 0.0001, *p=0.0131, **p=0.0055.

### Src kinase Blk is required for IL-23-induced mTORC1 activation and IL-17 production in γδ17 T cells

Dasatinib and Bosutinib are dual Src/Abl inhibitors. To identify their target in γδ17 T cells, we compared expression levels of all members of Abl and Src families using RNA-seq data available from the Immunological Genome Project (IGP) (Heng, Painter, and Immunological Genome Project Consortium, 2008). We observed that γδ17 T cells expressed minimal levels of the Abl kinases, whereas the B lymphoid kinase (Blk) was the most abundantly expressed Src kinase in these cells (**Fig. 7a**). Interestingly, Blk has previously been implicated in γδ17 T cell differentiation in the thymus (Laird, Laky and Hayes, 2010). To investigate the role of Blk in the IL-23 signalling cascade, we aimed to silence Blk expression using short hairpin RNAs (shBlk). shBlk was delivered to γδ17 T-LTIC cells via lentiviral constructs containing a GFP reporter. Upon transduction of γδ17 T-LTICs, we stimulated the cells with IL-23 and evaluated the effects of Blk silencing on mTORC1 activation and IL-17 production. We hypothesized that a reduction in pSrc-Y416 levels in shBlk-transduced cells would indicate that Blk was the major Src kinase member in these cells. Data in **Figure 7b** show a 60% decrease in pSrc-Y416 staining in LV-GFP-shBlk-transduced cells, suggesting that Blk is the major Src kinase in γδ17 T-LTICs. Next, we analysed pS6-S235/236 levels as a marker of mTORC1 activation. Our results revealed that IL-23-induced mTORC1 activation was significantly reduced in LV-GFP-shBlk-transduced cells (**Fig. 7c**). Finally, we examined cytokine production in response to IL-23 stimulation and found that IL-17A production was severely impaired in LV-GFP-shBlk-transduced cells (**Fig. 7d**). In conclusion, these experiments indicate that Blk is essential for IL-23-induced IL-17 production, likely through its role in enabling mTORC1 activation. These findings uncover a novel IL-23-Src-mTOR axis in type 3 immunity, and open the door for therapeutic targeting of Src signalling networks in IL-23/IL-17-driven inflammation.

**Figure 7.**
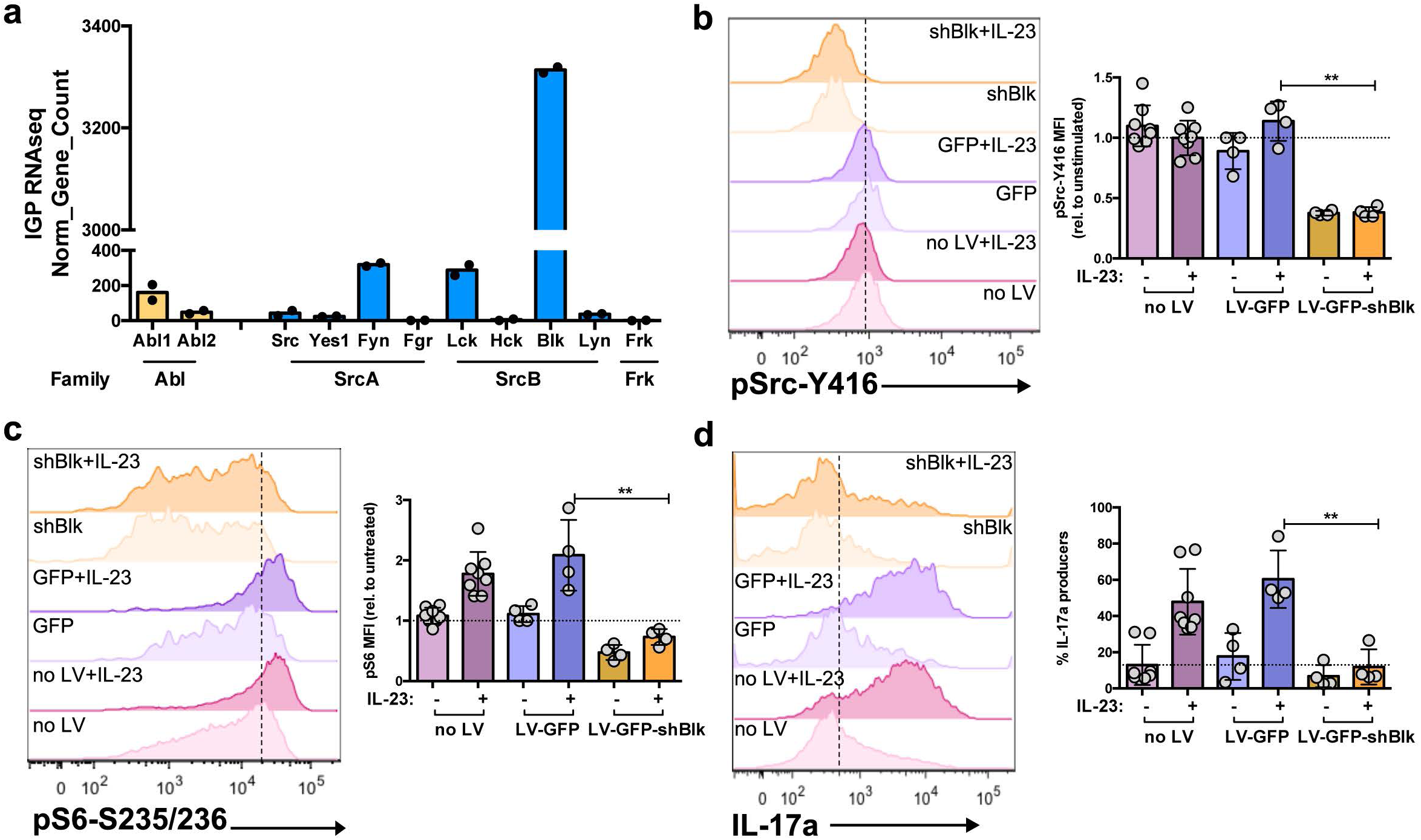
Silencing of the Src kinase Blk blocks mTORC1 activation and IL-17 production in response to IL-23. **a)** Graph shows normalized gene counts for all members of Src and Abl families in γδ17 T cells (mean, n=2), obtained from the Immunological Genome Project database (IGP). **b, c, d**) γδ17T-LTICs were transduced with lentiviral particles encoding GFP (LV-GFP) or GFP and a shRNA against Blk (LV-GFP-shBlk). 3-5 days later, cells were stimulated with IL-23 for 20h prior to flow cytometry analysis. **b**) Representative histograms show levels of phosphorylation of Src-Y416 residue (pSrc-Y416) in the different conditions. Graph show the mean fluorescence intensity (MFI), normalised to unstimulated condition (mean±sd, n=4-8). **c**) Representative histograms show levels of the indirect mTORC1 substrate pS6-S235/236 in the different conditions. Graph show the mean fluorescence intensity (MFI), normalised to unstimulated condition (mean±sd, n=4-8). **d)** Representative histograms show IL-17 production in the different conditions. Graph show the frequency of IL-17A producers (mean±sd, n=4-8). Statistical analysis was assessed using Kruskal-Wallis test with Dunńs correction for multiple comparisons. b) **p=0.0018, c)**p=0.0011, d) **p=0.0084.

## Discussion

The study of signalling networks that regulate the differentiation and effector function of type 3 immune cells has been hampered by the scarcity of IL-17-producing cells such as Th17 CD4 T cells, γδ17 T cells, and ILC3s. In this work, we developed a robust and scalable *in vitro* system based on long-term γδ17 T cell cultures (γδ17 T-LTICs), providing a faithful cellular model to study IL-23-driven IL-17 production. Using a drug-repurposing screening assay, we identified the dual Src/Abl kinase inhibitors Dasatinib and Bosutinib as potent suppressors of IL-23 signalling and IL-17 production *in vitro*. More importantly, Dasatinib attenuated psoriasis-like, IL-23-driven skin inflammation in the IMQ model when administered both systemically and topically. Our mechanistic studies revealed that Dasatinib and Bosutinib selectively block IL-23-induced mTORC1 and mTORC2 activation in γδ17 T-LTICs without affecting STAT3 phosphorylation. These results highlight a critical role for Src kinase family, particularly Blk, in coupling IL-23 signalling to mTORC1/mTORC2 activation and IL-17 production in γδ17 T cells. Our work underscores the value of γδ17 T-LTICs as a powerful platform to study IL-23 signalling and type 3 immune responses. Beyond IL-23-mediated effects, these cells also respond to IL-1β, which synergizes with IL-23 to induce proinflammatory mediators relevant in autoimmune diseases (Sutton *et al*., 2009; Coccia *et al*., 2012; Cai *et al*., 2014). Thus, γδ17 T-LTICs can contribute to future investigations of other regulatory mechanisms in these cells. Additionally, although γδ17 T-LTICs do not exhibit strong proliferative capacity, it is feasible to expand sufficient cell numbers for experimental approaches requiring medium-large amounts of starting material, such as proteomics, phosphoproteomics, or even western blotting. Thus, γδ17 T-LTICs represent a valuable and versatile tool for future research on γδ17 T cells and type 3 immune responses.

The data presented in this work demonstrate that Dasatinib administration exerts protective effects in an *in vivo* model of psoriasis-like skin inflammation. Both systemic and topical treatments reduced epidermal hyperproliferation, infiltration of CD4⁺ and γδ T cells, and the expansion of IL-17-producing cells. These findings suggest a potential repurposing of Dasatinib as an immunosuppressive agent in autoimmune diseases. Dasatinib is an orally available drug approved for the treatment of chronic myeloid leukemia and acute lymphoblastic leukemia, primarily due to its ability to inhibit Bcr-Abl activity. While it is also a known inhibitor of Src kinase family, well-known regulators of T cell signalling (Courtney, Lo and Weiss, 2018), its potential role as an immunosuppressant has not been extensively explored. Nevertheless, evidence from various mouse models of IL-23-related diseases supports the immunosuppressive properties of Dasatinib. For example, Dasatinib administration attenuated disease severity and delayed onset in the experimental autoimmune encephalomyelitis (EAE) model (Azizi *et al*., 2015). In rheumatoid arthritis models, Dasatinib alone (Guo *et al*., 2018; Min *et al*., 2023) or in combination with a subtherapeutic dose of anti-TNF (Ntari *et al*., 2021) showed therapeutic efficacy. In contrast, reports in inflammatory bowel diseases have described cases of Dasatinib-induced colitis in cancer patients (Del Sordo *et al*., 2022; Grillo *et al*., 2023; Gluch *et al*., 2025). These observations likely reflect the complex role of type 3 immune responses in the gut, where cytokines such as IL-23, IL-17, and IL-22 contribute to autoimmune inflammation but also play essential roles in maintaining barrier integrity (Mills, 2023). In this context, topical administration of Dasatinib may offer a promising strategy to limit systemic side effects. We show that topical application of Dasatinib effectively mitigates psoriasis-like skin inflammation, and in rheumatoid arthritis model a topical emulgel formulation containing Dasatinib has demonstrated protective effects (Donthi *et al*., 2023), supporting the feasibility of Dasatinib topical application. Together, these findings broaden our understanding of IL-23 signalling and support the potential of repurposing Src kinase inhibitors to treat IL-23/IL-17-driven inflammatory diseases. The efficacy of topical Dasatinib underscores its translational potential for skin-limited pathologies such as psoriasis and offers an alternative or complementary therapeutic approach. Psoriasis has a number of associated comorbidities. Of note, about 30% of psoriasis patients are prone to develop psoriatic arthritis (Ritchlin, Colbert and Gladman, 2017). Other comorbidities include metabolic syndrome and cardiovascular disorders, and the social stigma associated to the visibility of the lesions is associated to an increased risk for anxiety and depression (Parisi *et al*., 2013). Thus, therapeutic strategies should aim to manage disease severity but also to reduce the presence of comorbidities (Armstrong *et al*., 2025). In the context of precision medicine, it would be of great interest to determine if the interference with the newly described IL-23/Src/mTOR/IL-17 axis has a protective role in the development of psoriasis associated comorbidities. In addition to its immunosuppressive properties, Dasatinib has recently emerged as a senolytic compound. When used in combination with quercetin, known as the D+Q treatment, it is widely employed to selectively eliminate senescent cells (Lorenzo, Torrance and Haynes, 2023). Research into cellular senescence and its impact on pathological conditions is rapidly growing. Long-term D+Q treatment has been shown to be safe and well tolerated (Novais *et al*., 2021), and evidence in the literature suggests beneficial effects in autoimmune conditions such as Crohn’s disease (Sangfuang *et al*., 2025) and rheumatoid arthritis (de Jong *et al*., 2025). The ability of Dasatinib to simultaneously target immunopathogenic pathways and cellular senescence positions it as a unique therapeutic candidate for a range of chronic inflammatory diseases.

Our work has uncovered a novel IL-23-Src-mTOR signalling axis that plays an important role in IL-23/IL-17–driven pathologies. The activation of Src kinase family is a complex process, as these kinases can exist in various primed states. Beyond phosphorylation status, their intracellular localization and proximity to specific substrates are also critical determinants of their activity (Ortiz *et al*., 2021; Minguet, Maus and Schamel, 2025). In our study, we did not observe consistent changes in phosphorylation of the Src-Y416 residue following IL-23 stimulation, and we did not assessed the intracellular localization of the Src kinase family. Therefore, we cannot conclude whether IL-23 directly or indirectly activates Src kinases in γδ17 T cells. Interestingly, a connection between cytokine signalling, Src kinases, and mTOR activation has been previously described downstream of IL-2 receptor in CD8⁺ cytotoxic T cells (Ross *et al*., 2016). In this study, the Src inhibitor PP2 prevented IL-2-induced phosphorylation of S6 at Ser235/236 (readout of mTORC1 activity), without affecting phosphorylation of STAT5A-Y694 or STAT5B-Y699. These findings suggest that Jak-dependent networks may cooperate with Src signalling to regulate mTORC1 activity in T cells. In addition to the molecular mechanisms linking IL-23 and Src activity, another open question is how Src kinases modulate mTORC1 and mTORC2 activation. Notably, Src has been shown to promote mTORC1 recruitment to the lysosomal surface by regulating the GTP/GDP-bound state of Rag GTPases in response to amino acids (Pal et al., 2018). Additionally, other studies have suggested that Src-mediated activation of mTORC1 may occur independently of Akt signalling (Vojtechová et al., 2008). mTORC2 activation has been also shown to be regulated by Src in focal adhesions (Thompson *et al*., 2013). Further investigation into the precise molecular links between IL-23, Src kinase family, and mTOR complexes within the context of IL-17 biology could lead to novel therapeutic strategies to disrupt this pathogenic signalling network. Given the key role of mTOR in metabolic programming, targeting the IL-23-Src-mTOR axis could also modulate the broader transcriptional and epigenetic landscape of pathogenic effector cells(Patel and Powell, 2025).

Finally, our study identifies the Src kinase Blk as a critical mediator linking IL-23 signalling to mTORC1 and mTORC2 activation, thereby promoting IL-17 production in γδ17 T cells. While Src kinase family such as Lck and Fyn are well-established as early transducers of TCR signalling in conventional αβ T cells (Courtney, Lo and Weiss, 2018), their roles in unconventional T cell subsets have remained largely uncharacterized. In γδ17 T cells, Blk is the most abundantly expressed Src kinase and has previously been implicated in thymic differentiation of this subpopulation (Laird, Laky and Hayes, 2010). However, its function in peripheral effector responses was unknown. Our findings show that genetic silencing of Blk diminishes mTORC1 activity and impairs IL-17 production in γδ17 T cells, underscoring its essential role in IL-23-driven pathogenic responses. These results highlight the importance of Src kinases not only downstream of the TCR complex, but also reveal their critical function in cytokine signalling. We also observed that Dasatinib treatment affected CD4⁺ T cells *in vivo*, and inhibited Th17 cell differentiation in vitro. In these cells, the effects are likely mediated by Lck and/or Fyn, the predominant Src kinase in αβ T cells. Since TCR engagement by cognate antigen is essential for effector T cell differentiation, and Src kinases are key mediators of TCR signalling, it remains challenging to distinguish whether the effects of Src inhibition are due to interference with TCR signalling, IL-23 signalling, or more likely, both. Future work should aim to delineate how Src kinases are activated downstream of the IL-23 receptor in γδ17 T cells, Th17 cells and other IL-17-producing populations, which could inform new therapeutic strategies targeting IL-23–driven inflammation.

In summary, our study uncovers the Src kinase Blk as a novel signalling participant in the IL-23-mTOR-IL-17 axis in γδ17 T cells, and supports the therapeutic potential of Src inhibitors in IL-23/IL-17–mediated diseases. Notably, Dasatinib, a clinically approved Src/Abl inhibitor, emerges as a promising candidate for repurposing in autoimmune and inflammatory conditions. Its combined immunomodulatory and senolytic effects, along with the potential for topical delivery, warrant further clinical investigation.

## Acknowledges

This work was supported by grants from the Spanish Goverment to MNN (Ministry of Science and Innovation PID2019-110511RB-I00 and Ministry of Science, Innovation and Universities, PID2022-141729OB-I00). Institutional grant from the Fundación Ramón Areces to the CBM is also acknowledged. The funders had no role in study design, data collection and analysis, decision to publish, or preparation of the manuscript. We acknowledge Dr. Maria L. Toribio, Dr. Maria Mittelbrunn and their groups for advice, assistance, and critical discussion. We also thank Dr. Jose Cuezva for providing the FDA-approved drug library. We thank the Histology Facility at CNB-CSIC for the histological preparation of biological samples.

## Author contribution

MJV, IC, JSY, LS-R and GP-F Investigation, Formal analysis, Validation, Writing –review & editing.

AO and JT Investigation, Validation.

MNN Conceptualization, Data curation, Formal analysis, Funding acquisition, Investigation, Methodology, Validation, Writing - original draft, Writing –review & editing.

## Declaration of Generative AI and AI-assisted technologies in the writing process

During the preparation of this work the author(s) used ChatGPT in order to improve readability and language. After using this tool/service, the author(s) reviewed and edited the content as needed and take(s) full responsibility for the content of the publication

---

**Supplemental Figure 1.**
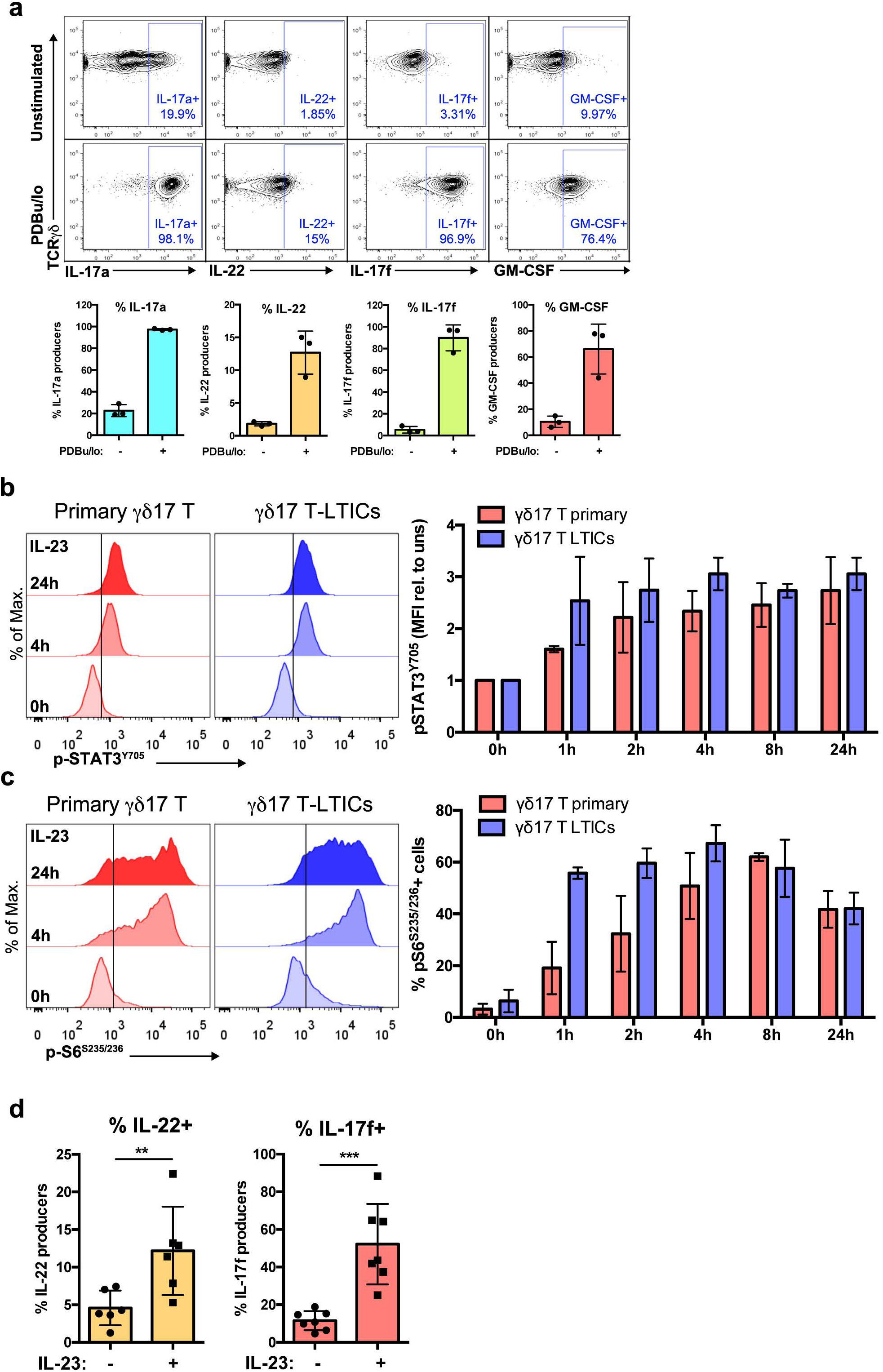
Characterisation of γδ17T-LTICs cells. **a)** γδ17T-LTICs were stimulated with phorbol 12,13-dibutyrate (PDBu) and Ionomycin (Io) for 4h in presence of Golgi-Plug, and processed for flow cytometry analysis. Representative dot plots show TCRγδ expression against IL-17A, IL-22, IL-17f and GM-CSF production. Graphs represent the frequency of cytokine-producing cells (mean±sd, n=3 independent cultures). **b)** Primary γδ17T cells (isolated and expanded in IL-7 for 5 days) and γδ17T -LTICs were stimulated with IL-23 for different lengths of time, and processed for analysis of STAT3-Y705 phosphorylation. Representative histograms show p-STAT3-Y705 at the indicated time points. Graph show the mean fluorescence intensity (MFI), relative to unstimulated condition (uns) (mean±sd, n=3 independent experiments). **c)** Cells were treated as in b), and S6-S235/236 phosphorylation was assessed by flow cytometry. **d)** γδ17T-LTICs were stimulated with IL-23 for 20h, and Golgi-Plug was added different the last 4h of incubation, followed by analysis by flow cytometry. Graphs show the frequency of IL-22 and IL-17f-producing cells (mean±sd, n=6 independent cultures).

**Supplemental Figure 2.**
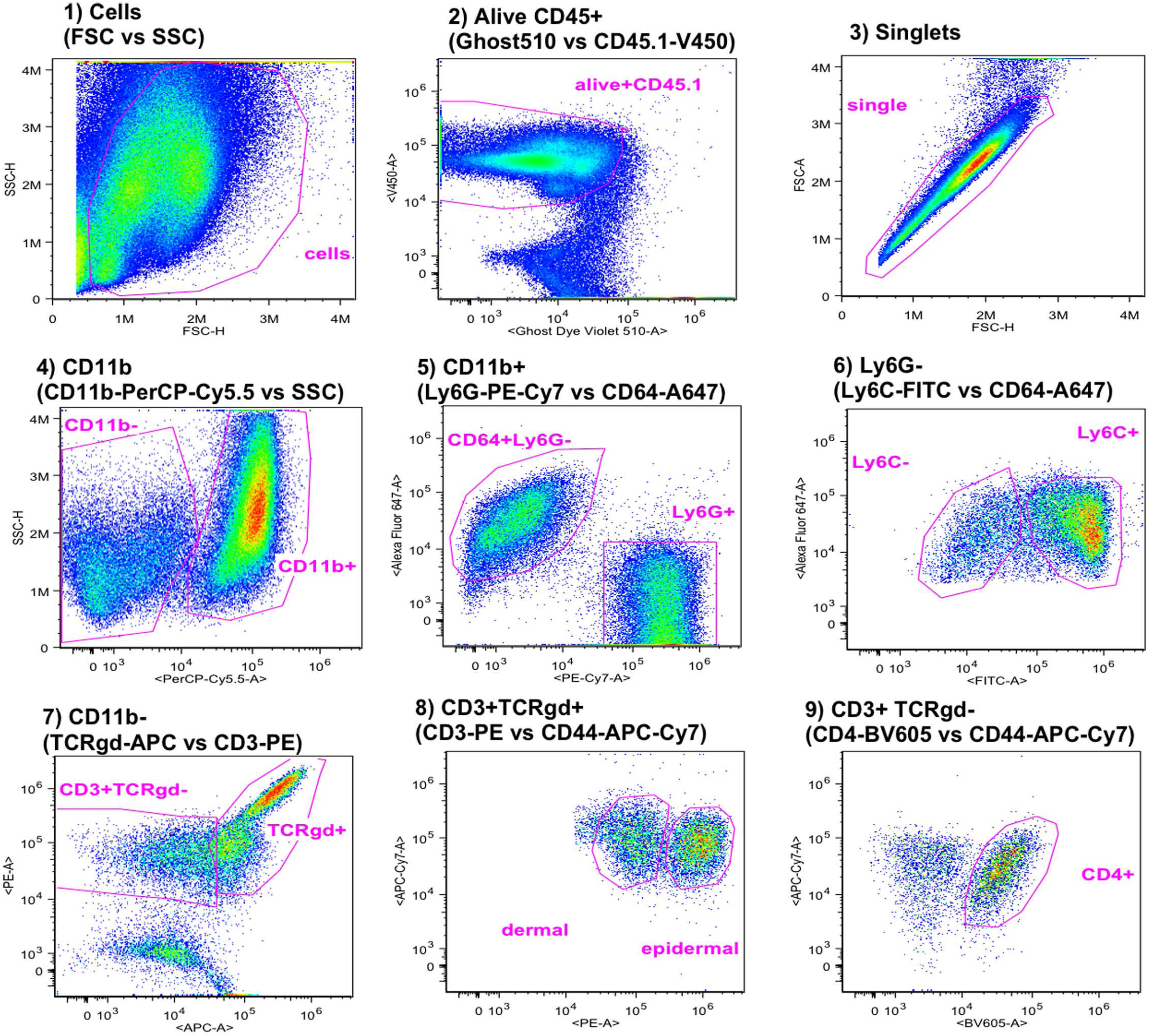
Gating strategy for analysis of skin infiltrate in Imiquimod-induced skin inflammation model. Ears were minced and subjected to enzymatic digestion prior to flow cytometry analysis. Representative dot plots show the gating strategy for analysis of leucocyte subpopulations infiltrating the ear skin. **(1)** Cells were gated according to their FSC-H and SSC-H profile, **(2)** alive leucocytes were gated as CD45^+^ and Ghost-Dye^neg^, **(3)** single cells were gated according to their FSC-H and FSC-A profile, **(4)** cells were gated as CD11b^+^ (myeloid) or CD11b^neg^ (lymphoid), **(5)** Myeloid cells (CD11b^+^) were gated as CD64^neg^Ly6G^+^ (neutrophils) and CD64^+^Ly6G^neg^, **(6)** CD64^+^Ly6G^neg^ were further subdivided into Ly6C^+^ (inflammatory monocyte-macrophage) and Ly6C^neg^ (resident monocyte). **(7)** Lymphoid cells (CD11b^neg^) were defined as CD3^+^TCRγδ^+^ and CD3^+^TCRγδ^neg^. **(8)** TCRγδ^+^ cells were further divided into dermal and epidermal subpopulations based on TCRγδ expression level (TCRγδ^int^ and TCRγδ^hi^, respectively). **(9)** CD3^+^TCRγδ^neg^ cells were further characterised based on CD4 expression.

**Supplemental Table. Drug screening raw data** γδ17 T-LTICs were stimulated with IL-23 in presence of 88 FDA-approved drugs for 20h in three independent biological replicates. IL-17A production was determined by ELISA, and cell viability was measured by flow cytometry. **Tab1**) L1300-01 plate layout. Experimental design of the 96 well plates used for screening. Each well specifies the assayed FDA-approved drug. **Tab2**) L1300-01 FDA-approved drug information. Columns specify Selleckem Catalog number, Item name, Plate location (as shown in Tab1), molecular weight (M.w.), CAS number, Indication, Target and Brief Description of each drug. Tab3) Screening results. Table provide the individual screening results (IL-17 production and viability, normalised to the IL-23 condition), for the three biological replicates (BR1, BR2, BR3).

## Bibliography

Annunziato, F., Romagnani, C. and Romagnani, S. (2015) ‘The 3 major types of innate and adaptive cell-mediated effector immunity’, The Journal of Allergy and Clinical Immunology, 135(3), pp. 626–635. Available at: 10.1016/j.jaci.2014.11.001.

Armstrong, A.W. et al. (2025) ‘Psoriasis’, Nature Reviews. Disease Primers, 11(1), p. 45. Available at: 10.1038/s41572-025-00630-5.

Awasthi, A. et al. (2009) ‘Cutting edge: IL-23 receptor gfp reporter mice reveal distinct populations of IL-17-producing cells.’, Journal of immunology (Baltimore, Md. : 1950), 182(10), pp. 5904–5908. Available at: 10.4049/jimmunol.0900732.

Azizi, G. et al. (2015) ‘Therapeutic effects of dasatinib in mouse model of multiple sclerosis’, Immunopharmacology and Immunotoxicology, 37(3), pp. 287–294. Available at: 10.3109/08923973.2015.1028074.

Cai, Y. et al. (2011) ‘Pivotal role of dermal IL-17-producing γδ T cells in skin inflammation.’, Immunity, 35(4), pp. 596–610. Available at: https://doi.org/©.

Cai, Y. et al. (2014) ‘Differential developmental requirement and peripheral regulation for dermal Vγ4 and Vγ6T17 cells in health and inflammation.’, Nature communications, 5, p. 3986. Available at: 10.1038/ncomms4986.

Cai, Y. et al. (2019) ‘Differential Roles of the mTOR-STAT3 Signaling in Dermal γδ T Cell Effector Function in Skin Inflammation.’, Cell reports, 27(10), pp. 3034–3048.e5. Available at: 10.1016/j.celrep.2019.05.019.

Chan, J.R. et al. (2006) ‘IL-23 stimulates epidermal hyperplasia via TNF and IL-20R2-dependent mechanisms with implications for psoriasis pathogenesis.’, The Journal of experimental medicine, 203(12), pp. 2577–2587. Available at: 10.1084/jem.20060244.

Cibrian, D. et al. (2020) ‘Targeting L-type amino acid transporter 1 in innate and adaptive T cells efficiently controls skin inflammation.’, The Journal of allergy and clinical immunology, 145(1), pp. 199–214.e11. Available at: 10.1016/j.jaci.2019.09.025.

Coccia, M. et al. (2012) ‘IL-1β mediates chronic intestinal inflammation by promoting the accumulation of IL-17A secreting innate lymphoid cells and CD4(+) Th17 cells.’, The Journal of experimental medicine, 209(9), pp. 1595–1609. Available at: 10.1084/jem.20111453.

Courtney, A.H., Lo, W.-L. and Weiss, A. (2018) ‘TCR Signaling: Mechanisms of Initiation and Propagation’, Trends in Biochemical Sciences, 43(2), pp. 108–123. Available at: 10.1016/j.tibs.2017.11.008.

Del Sordo, R. et al. (2022) ‘Imatinib and Dasatinib-induced Ulcerative Colitis: Case Report’, Inflammatory Bowel Diseases, 28(1), pp. e1–e2. Available at: 10.1093/ibd/izab196.

Donthi, M.R. et al. (2023) ‘Dasatinib-Loaded Topical Nano-Emulgel for Rheumatoid Arthritis: Formulation Design and Optimization by QbD, In Vitro, Ex Vivo, and In Vivo Evaluation’, Pharmaceutics, 15(3), p. 736. Available at: 10.3390/pharmaceutics15030736.

Fiala, G.J., Gomes, A.Q. and Silva-Santos, B. (2020) ‘From thymus to periphery: Molecular basis of effector γδ-T cell differentiation’, Immunological Reviews, 298(1), pp. 47–60. Available at: 10.1111/imr.12918.

van der Fits, L. et al. (2009) ‘Imiquimod-induced psoriasis-like skin inflammation in mice is mediated via the IL-23/IL-17 axis.’, Journal of immunology (Baltimore, Md. : 1950), 182(9), pp. 5836–5845. Available at: 10.4049/jimmunol.0802999.

Gluch, A.E. et al. (2025) ‘Dasatinib-induced colitis in a patient with chronic myeloid leukaemia’, BMJ case reports, 18(1), p. e263646. Available at: 10.1136/bcr-2024-263646.

Gray, E.E. et al. (2013) ‘Deficiency in IL-17-committed Vγ4(+) γδ T cells in a spontaneous Sox13-mutant CD45.1(+) congenic mouse substrain provides protection from dermatitis’, Nature Immunology, 14(6), pp. 584–592. Available at: 10.1038/ni.2585.

Grillo, F. et al. (2023) ‘Dasatinib-induced Crohn’s-like colitis’, Journal of Clinical Pathology, 76(3), pp. 202–205. Available at: 10.1136/jclinpath-2022-208340.

Guo, K. et al. (2018) ‘Treatment Effects of the Second-Generation Tyrosine Kinase Inhibitor Dasatinib on Autoimmune Arthritis’, Frontiers in Immunology, 9, p. 3133. Available at: 10.3389/fimmu.2018.03133.

Hartwig, T. et al. (2018) ‘Regulatory T Cells Restrain Pathogenic T Helper Cells during Skin Inflammation’, Cell Reports, 25(13), pp. 3564–3572.e4. Available at: 10.1016/j.celrep.2018.12.012.

Heng, T.S.P., Painter, M.W., and Immunological Genome Project Consortium (2008)‘ The Immunological Genome Project: networks of gene expression in immune cells.’, Nature immunology, 9(10), pp. 1091–1094. Available at: 10.1038/ni1008-1091.

Hwang, J.-R. et al. (2020) ‘Recent insights of T cell receptor-mediated signaling pathways for T cell activation and development’, Experimental & Molecular Medicine, 52(5), pp. 750–761. Available at: 10.1038/s12276-020-0435-8.

Inácio, D. et al. (2025) ‘Signature cytokine-associated transcriptome analysis of effector γδ T cells identifies subset-specific regulators of peripheral activation’, Nature Immunology [Preprint]. Available at: 10.1038/s41590-024-02073-8.

Ivanov, I.I. et al. (2006) ‘The orphan nuclear receptor RORgammat directs the differentiation program of proinflammatory IL-17+ T helper cells.’, Cell, 126(6), pp. 1121–1133. Available at: 10.1016/j.cell.2006.07.035.

de Jong, T.A. et al. (2025) ‘Senolytic treatment to rescue hallmarks of senescence in lymph node fibroblasts from patients with rheumatoid arthritis: Implications for premature aging and potential therapeutic intervention in early rheumatoid arthritis’, Clinical and Experimental Immunology, 219(1), p. uxaf029. Available at: 10.1093/cei/uxaf029.

Laird, R.M., Laky, K. and Hayes, S.M. (2010) ‘Unexpected role for the B cell-specific Src family kinase B lymphoid kinase in the development of IL-17-producing γδ T cells.’, Journal of immunology (Baltimore, Md. : 1950), 185(11), pp. 6518–6527. Available at: 10.4049/jimmunol.1002766.

Lorenzo, E.C., Torrance, B.L. and Haynes, L. (2023) ‘Impact of senolytic treatment on immunity, aging, and disease’, Frontiers in Aging, 4, p. 1161799. Available at: 10.3389/fragi.2023.1161799.

Mezghiche, I. et al. (2024) ‘Interleukin 23 receptor: Expression and regulation in immune cells’, European Journal of Immunology, 54(1), p. 2250348. Available at: 10.1002/eji.202250348.

Michel, M.-L. et al. (2012) ‘Interleukin 7 (IL-7) selectively promotes mouse and human IL-17-producing γδ cells.’, Proceedings of the National Academy of Sciences of the United States of America, 109(43), pp. 17549–17554. Available at: 10.1073/pnas.1204327109.

Mills, K.H.G. (2023) ‘IL-17 and IL-17-producing cells in protection versus pathology’, Nature Reviews. Immunology, 23(1), pp. 38–54. Available at: 10.1038/s41577-022-00746-9.

Min, H.K. et al. (2023) ‘Dasatinib, a selective tyrosine kinase inhibitor, prevents joint destruction in rheumatoid arthritis animal model’, International Journal of Rheumatic Diseases, 26(4), pp. 718–726. Available at: 10.1111/1756-185X.14627.

Minguet, S., Maus, M.V. and Schamel, W.W. (2025) ‘From TCR fundamental research to innovative chimeric antigen receptor design’, Nature Reviews Immunology, 25(3), pp. 212–224. Available at: 10.1038/s41577-024-01093-7.

Muschaweckh, A., Petermann, F. and Korn, T. (2017) ‘IL-1β and IL-23 Promote Extrathymic Commitment of CD27+CD122-γδ T Cells to γδT17 Cells.’, Journal of immunology (Baltimore, Md. : 1950), 199(8), pp. 2668–2679. Available at: 10.4049/jimmunol.1700287.

Novais, E.J. et al. (2021) ‘Long-term treatment with senolytic drugs Dasatinib and Quercetin ameliorates age-dependent intervertebral disc degeneration in mice’, Nature Communications, 12(1). Available at: 10.1038/s41467-021-25453-2.

Ntari, L. et al. (2021) ‘Combination of subtherapeutic anti-TNF dose with dasatinib restores clinical and molecular arthritogenic profiles better than standard anti-TNF treatment’, Journal of Translational Medicine, 19(1), p. 165. Available at: 10.1186/s12967-021-02764-y.

Ortiz, M.A. et al. (2021) ‘Src family kinases, adaptor proteins and the actin cytoskeleton in epithelial-to-mesenchymal transition’, Cell Communication and Signaling, 19(1). Available at: 10.1186/s12964-021-00750-x.

Pantelyushin, S. et al. (2012) ‘Rorγt+ innate lymphocytes and γδ T cells initiate psoriasiform plaque formation in mice.’, The Journal of clinical investigation, 122(6), pp. 2252–2256. Available at: 10.1172/JCI61862.

Papotto, P.H. et al. (2017) ‘IL-23 drives differentiation of peripheral γδ17 T cells from adult bone marrow-derived precursors.’, EMBO reports, 18(11), pp. 1957–1967. Available at: 10.15252/embr.201744200.

Parham, C. et al. (2002) ‘A receptor for the heterodimeric cytokine IL-23 is composed of IL-12Rbeta1 and a novel cytokine receptor subunit, IL-23R.’, Journal of immunology (Baltimore, Md. : 1950), 168(11), pp. 5699–5708.

Parisi, R. et al. (2013) ‘Global epidemiology of psoriasis: a systematic review of incidence and prevalence.’, The Journal of investigative dermatology, 133(2), pp. 377–385. Available at: 10.1038/jid.2012.339.

Patel, C.H. and Powell, J.D. (2025) ‘More TOR: The expanding role of mTOR in regulating immune responses’, Immunity, 58(7), pp. 1629–1645. Available at: 10.1016/j.immuni.2025.06.010.

Ramírez-Valle, F., Gray, E.E. and Cyster, J.G. (2015) ‘Inflammation induces dermal Vγ4+ γδT17 memory-like cells that travel to distant skin and accelerate secondary IL-17-driven responses.’, Proceedings of the National Academy of Sciences of the United States of America, 112(26), pp. 8046–8051. Available at: 10.1073/pnas.1508990112.

Reid, C. and Griffiths, C.E.M. (2020) ‘Psoriasis and Treatment: Past, Present and Future Aspects’, Acta Dermato-Venereologica, 100(3), p. adv00032. Available at: 10.2340/00015555-3386.

Ritchlin, C.T., Colbert, R.A. and Gladman, D.D. (2017) ‘Psoriatic Arthritis’, New England Journal of Medicine. Edited by D.L. Longo, 376(10), pp. 957–970. Available at: 10.1056/nejmra1505557.

Ross, S.H. et al. (2016) ‘Phosphoproteomic Analyses of Interleukin 2 Signaling Reveal Integrated JAK Kinase-Dependent and -Independent Networks in CD8(+) T Cells.’, Immunity, 45(3), pp. 685–700. Available at: 10.1016/j.immuni.2016.07.022.

Sangfuang, N. et al. (2025) ‘Effects of senotherapeutics on gut microbiome dysbiosis and intestinal inflammation in Crohn’s disease: A pilot study’, Translational Research: The Journal of Laboratory and Clinical Medicine, 278, pp. 36–47. Available at: 10.1016/j.trsl.2025.02.004.

Schmolka, N. et al. (2013) ‘Epigenetic and transcriptional signatures of stable versus plastic differentiation of proinflammatory γδ T cell subsets.’, Nature immunology, 14(10), pp. 1093–1100. Available at: 10.1038/ni.2702.

Sutton, C.E. et al. (2009) ‘Interleukin-1 and IL-23 induce innate IL-17 production from gammadelta T cells, amplifying Th17 responses and autoimmunity.’, Immunity, 31(2), pp. 331–341. Available at: 10.1016/j.immuni.2009.08.001.

Thompson, W.R. et al. (2013) ‘Mechanically activated Fyn utilizes mTORC2 to regulate RhoA and adipogenesis in mesenchymal stem cells’, Stem Cells (Dayton, Ohio), 31(11), pp. 2528–2537. Available at: 10.1002/stem.1476.

Yang, Y. et al. (2014) ‘Focused specificity of intestinal TH17 cells towards commensal bacterial antigens.’, Nature, 510(7503), pp. 152–156. Available at: 10.1038/nature13279.

Yoshiki, R. et al. (2014) ‘IL-23 from Langerhans cells is required for the development of imiquimod-induced psoriasis-like dermatitis by induction of IL-17A-producing γδ T cells.’, The Journal of investigative dermatology, 134(7), pp. 1912–1921. Available at: 10.1038/jid.2014.98.

Zheng, Y. et al. (2007) ‘Interleukin-22, a T(H)17 cytokine, mediates IL-23-induced dermal inflammation and acanthosis.’, Nature, 445(7128), pp. 648–651. Available at: 10.1038/nature05505.

